# Epidermal basal domains organization highlights skin robustness to environmental exposure

**DOI:** 10.1101/2022.02.23.481662

**Authors:** Sangeeta Ghuwalewala, Seon A Lee, Kevin Jiang, Joydeep Baidya, Gopal Chovatiya, Pritinder Kaur, David Shalloway, Tudorita Tumbar

## Abstract

Adult interfollicular epidermis (IFE) renewal is likely orchestrated by physiological demands of its complex tissue architecture comprising spatial and cellular heterogeneity. Mouse tail and back skin display two kinds of basal IFE spatial domains that regenerate at different rates. Here we elucidate the molecular and cellular states of basal IFE domains by marker expression and single cell transcriptomics in mouse and human skin. We uncover two paths of basal cell differentiation that reflect in part the IFE spatial domain organization. We unravel previously unrecognized similarities between mouse tail IFE basal domains defined as scales and interscales versus human rete ridges and inter-ridges, respectively. Second, our basal IFE transcriptomics and gene targeting in mice provide evidence supporting a physiological role of IFE domains: adaptation to differential UV exposure. We identify Sox6 as a novel UV-induced and interscale/inter-ridge basal IFE-domain transcription factor, important for IFE proliferation and survival. The spatial, cellular, and molecular organization of IFE basal domains underscores skin adaptation to environmental exposure and its unusual robustness in adult homeostasis.

**Synopsis:** 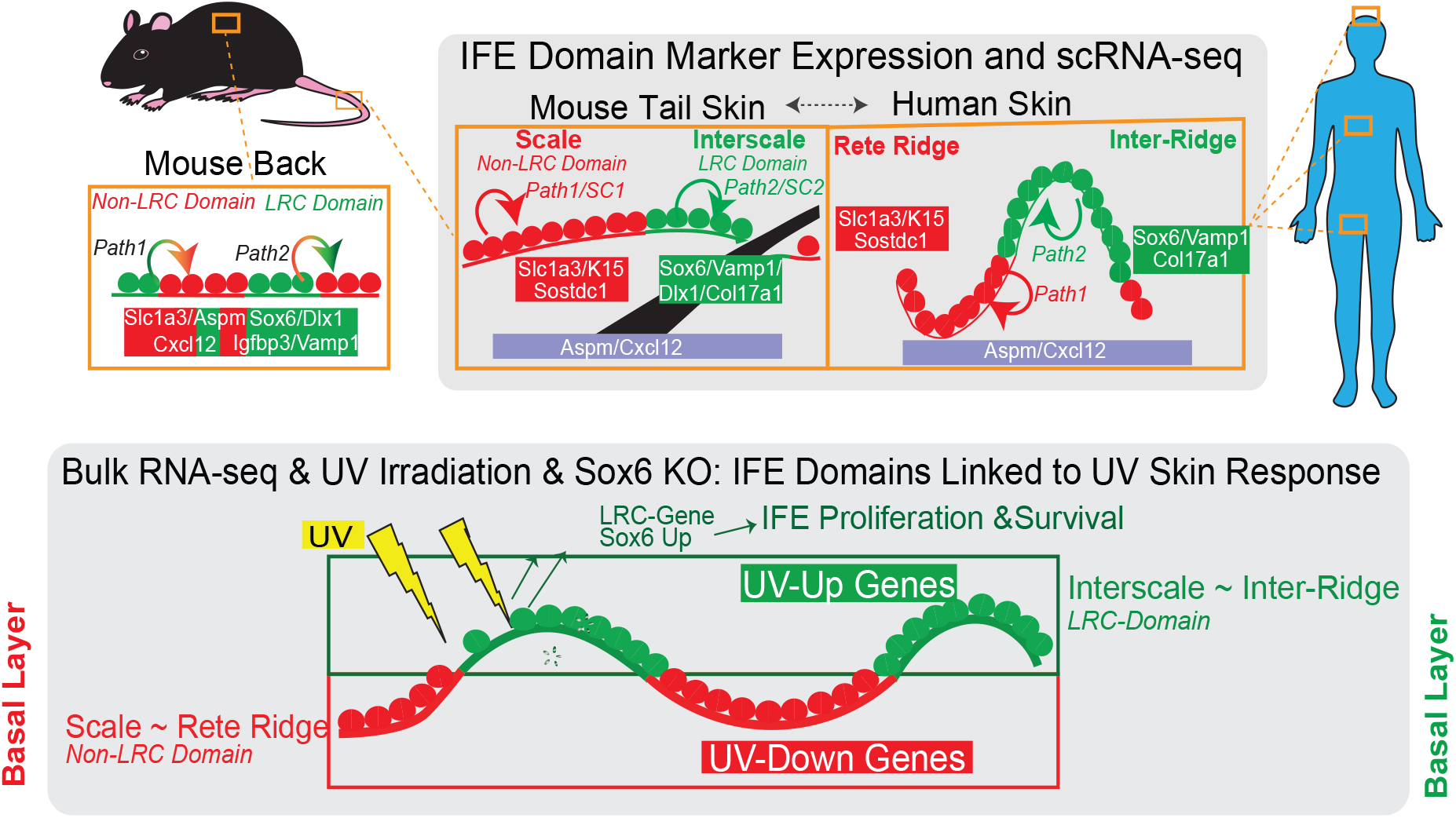

## Introduction

Adult skin interfollicular epidermis (IFE) makes the essential body barrier to the outside world and must respond to a variety of environmental insults and physiological demands (Blanpain & Fuchs, 2009; Gola & Fuchs, 2021). Homeostatic IFE renewal is likely resulting from local microenvironmental challenges, locally imposed on an intricate tissue architecture that comprises multiple levels of heterogeneity. The vicinity of the hair follicle, body region variations, skin maturation and aging, and stress conditions such as mechanical stretching or wounding pose additional challenges to IFE renewal (Aragona *et al*, 2017; Aragona *et al*, 2020; Ichijo *et al*, 2021; Park *et al*, 2017; Roy *et al*, 2016; Roy *et al*, 2020). Development of skin cancers, such as melanoma, are also associated with IFE regional heterogeneity and UV-exposure (Kohler *et al*, 2017; Moon *et al*, 2017). We and others have previously documented that adult mouse IFE presents two spatially distinct domains that renew at different rates during homeostasis. These two domains are found in both mouse tail skin, known as scales and interscales, and in back skin; they can be identified as H2B-GFP or BrdU non-label retaining cells (non-LRCs) domains and LRCs domains (Aragona *et al*., 2017; Gomez *et al*, 2013; Riquelme *et al*, 2008; Roy *et al*., 2016; Sada *et al*, 2016; Sanchez-Danes *et al*, 2016). Human skin also presents two kinds of spatial domains, known as rete ridges and inter-ridge (Lawlor & Kaur, 2015), which based on their location in the epidermis have differential exposure to the outside environment. The functional significance of the two spatial IFE domains, and their correspondence from mouse to human skin are currently unknown. Recent single-cell transcriptomics of mouse skin revealed basal cell heterogeneity with multiple cell states (Aragona *et al*., 2020; Dekoninck *et al*, 2020; Haensel *et al*, 2020; Joost *et al*, 2018; Lin *et al*, 2020), but how these states are spatially and functionally organized in the IFE is unclear. Recent data from human newborn foreskin suggests that different basal IFE cell states might be spatially organized (Wang *et al*, 2020). A persistent lack of markers to distinguish spatial and functional basal cellular subsets impedes our current understanding of IFE organization.

Here, we use markers of basal IFE cellular subsets in mouse and human skin to examine the molecular, cellular, and functional organization of two IFE spatial domains. We show that the two IFE domains have different gene expression patterns and predominantly different paths of basal layer cell differentiation (Synopsis). Despite these differences, they both contain comparable mixtures of cell states previously defined as stem, proliferating, and differentiating basal cell types. We find that mouse back skin IFE domain organization stands out as somewhat unique, but we unearth previously un-recognized similarities of basal IFE organization between mouse tail scales and interscales with human rete ridges and inter-ridges respectively (Synopsis). Different molecular pathways—notably UV-response genes— are differentially upregulated in the two IFE domains, corresponding to their differential exposure to the outside environment. Our gene targeting functional studies in mice support the hypothesis that this heterogeneity is physiologically important, enabling the skin’s adaptive response to UV exposure (Synopsis). This spatial and molecular organization of the IFE may help explain the remarkable robustness of long-term skin homeostasis in the face of environmental challenges.

## Results

### Mouse tail scale/non-LRC vs interscale/LRC basal IFE domains express different levels of markers

We previously labeled skin cells based on proliferation history and isolated IFE label retaining cells (LRCs) and non-LRCs from mouse back skin using our tet-repressible K5tTA x pTRE-H2BGFP transgenic mice (Sada *et al*., 2016; Tumbar *et al*, 2004). A classical stem cell (SC)-transit amplifying (TA) cell model (Kaur & Potten, 2011), predicted that LRCs would be exclusively SCs whereas non-LRCs were TA cells. Instead, we found that both LRCs and non-LRCs contained long-lived regenerative (stem) cells, which could be marked with distinct genetic drivers (Dlx1-CreER and Slc1a3-CreER respectively). Moreover, LRCs and non-LRCs and their corresponding marked lineages segregated in two spatial domains in mouse back and tail skin (Figure 1A and (Sada *et al*., 2016)). In tail skin, LRC domains correspond to interscales, which include a sub-region uniting the hair follicle that we call the ‘line’, whereas non-LRC domains corresponded to scales (Figure 1A, S1A,B and (Sada *et al*., 2016)), findings also reported by other groups (Dekoninck *et al*., 2020; Gomez *et al*., 2013; Sada *et al*., 2016; Sanchez-Danes *et al*., 2016). However, two other groups reported a lack of IFE LRCs after 1-week chase (Piedrafita *et al*, 2020; Rompolas *et al*, 2016). Using two inducible systems (K14-rTA and K5-tTA) we found that 2-week chase via K5-tTA renders easily detectable LRCs. This occurs not only in tail interscales and back skin, as expected (Figure S1C) (Sada *et al*., 2016), but also in other body regions such as ear and paw (Figure S2). We also used tamoxifen (TM) induction experiments and confirmed preferential Slc1a3-CreER or Dlx1-CreER marking of non-LRCs or LRCs spatial clusters in tail (Figure 1B, C), back, ear and paw skin (Figure S2A-D). Next, we tested marking via Aspm-CreER, another basal non-LRC gene from the back skin microarray (Kang *et al*, 2020; Sada *et al*., 2016), but this was not enriched in scale (non-LRC) domain (Figure 1B, C and S1A). Interestingly, the Dlx1-CreER marking was enriched along the interscale/scale boundary and all three drivers marked the interscale ‘line’ sub-structure (Figure 1A-C and (Sada *et al*., 2016)). This data suggests that although the H2B-GFP non-LRCs cluster preferentially in scales and LRCs in interscales, the LRC/non-LRC gene expression spatial patterns are more complex.

**Figure 1:**
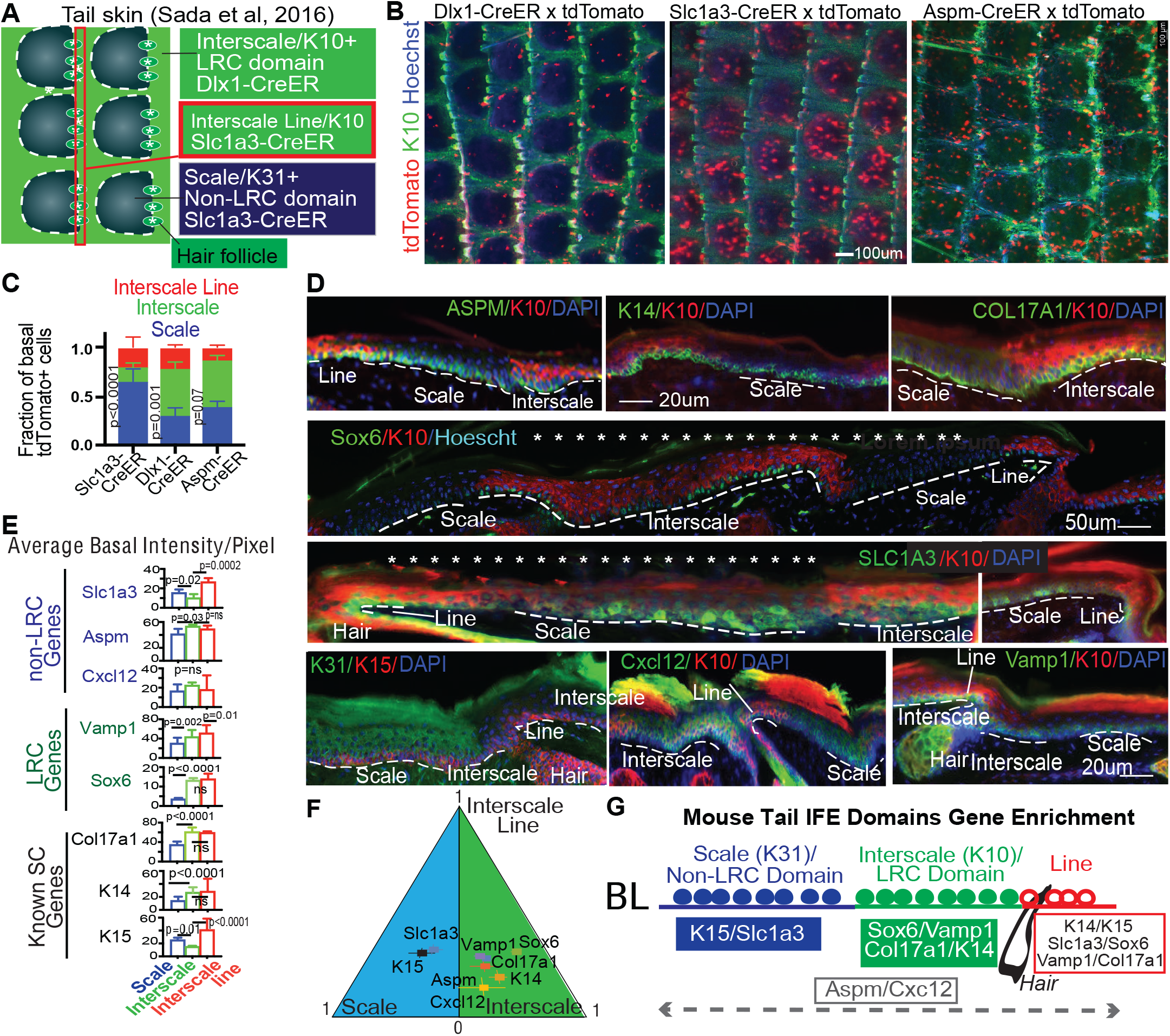
Basal layer cells express different LRC/non-LRC markers in scales vs interscales of mouse tail skin. (A) Model of tail skin organization in IFE domains. (B) Whole mount low magnification images show marking in interscale (K10+) vs scale regions, quantified in (C). (D) Tail skin sections immunofluorescence stained for interscale (K10) or scale (K31) supra-basal markers, and basal markers. (E) Quantification of images like those in (d) with n=3-4 mice and 6-7 images per mouse showing marker enrichment. P-values calculated by Student’s t-test (mixed effect model). (F) Relative gene expression/area levels from (D) and (E), normalized to sum to one, and displayed in a barycentric plot. Error bars are SEMs. (G) Cartoon summarizing spatial distribution of markers in the tail IFE domains.

To examine spatial patterns of protein expression in the IFE basal domains, we used immunofluorescence (IF) staining and microscopy on different body regions of mice at different ages. We tested commercial antibodies to >10 gene candidates we previously identified by microarrays of back skin IFE LRCs/non-LRCs (Sada *et al*., 2016), and obtained working conditions for Sox6 and Vamp1 (enriched in LRCs) and Slc1a3, Cxcl12, and Aspm (enriched in non-LRCs). We also used antibodies to some previously known SC-enriched genes - K15 (Cotsarelis, 2006; Kaur, 2004), Col17a1(Li *et al*, 2019; Watanabe *et al*, 2017) and K14(Mascre *et al*, 2012). Most LRC/non-LRC enriched markers, but especially Slc1a3, were expressed in distinct basal cell clusters in adult tail (Figure 1D and S1D) and back, ear and paw (Figure S2E), but newborn skin was more homogeneous (Figure S3). Tail image quantification measured the differential expression of the markers in the scale, interscale, and line basal IFE regions (Figure 1E). Except for Aspm and Cxcl12, the non-LRC markers (e.g. Slc1a3) and LRC markers (Sox6, Vamp1) were enriched in scale vs. interscale, respectively (Figure 1F). Unexpectedly, previously known SC markers also showed preferred expression: K15 was enriched in scale and K14 and Col17a1 were enriched in interscale (Figure 1E, F).

In summary, we identified markers that distinguish the basal layer of scale (*K15/Slc1a3*) as a non-LRC tail domain from the basal layer of interscale (*Vamp1/Sox6/Col17a1/K14*) as a LRC tail domain (Figure 1G). These molecular differences between basal IFE spatial LRC/non-LRC domains also exist to some extent in other body regions of the mouse skin.

### Human inter-ridges vs rete ridges resemble the mouse interscale vs scale domains

Human IFE is characterized by an undulating pattern of rete ridges (RR), which are spatial domains embedded deep into the dermis, and inter-ridges (IR), which are more raised IFE spatial domains (Figure 2A and (Lawlor and Kaur, 2015)). Here we used humans skin samples to probe for expression of mouse IFE LRC domain (interscale in tail) and non-LRC domain (scale in tail) markers. Strikingly, we found preferential expression of Slc1a3/K15 in the rete ridges and of Vamp1, Sox6, and Col17a1 in the inter-ridges (Figures 2A-H and S4A-D). This correlation is maintained to some extent in human breast, forehead, abdomen, scalp, arm, cheek, ear, and newborn foreskin (ages 21, 30, 45, 52, 76-year-old; Figure S4B and Table S1). These data reveal previously unrecognized similarities between mouse tail and human IFE organization in basal layer. Specifically, mouse tail interscales/LRC domains correspond to inter-ridges in human skin, and scales/non-LRC domains correspond to rete ridges (compare Figure 1G and 2M). Other basal markers such as Cxcl12 and Aspm were found in small basal cell clusters in both IR and RR (Figure 2D, E), reminiscent of their presence in both scales and interscales in mouse tail (Figure 1D-G). We briefly inquired into the proliferative status of RR and IR, but Ki67^+^ proliferative cells showed no obvious difference (Figure S4A), as reported (Ruetze *et al*., 2010). Notably, marker expression heterogeneity in the newborn human foreskin was less pronounced than in aged skin (Figure S4C, D), reminiscent of the mouse age-dependent differences in heterogeneity (Figure S3).

**Figure 2:**
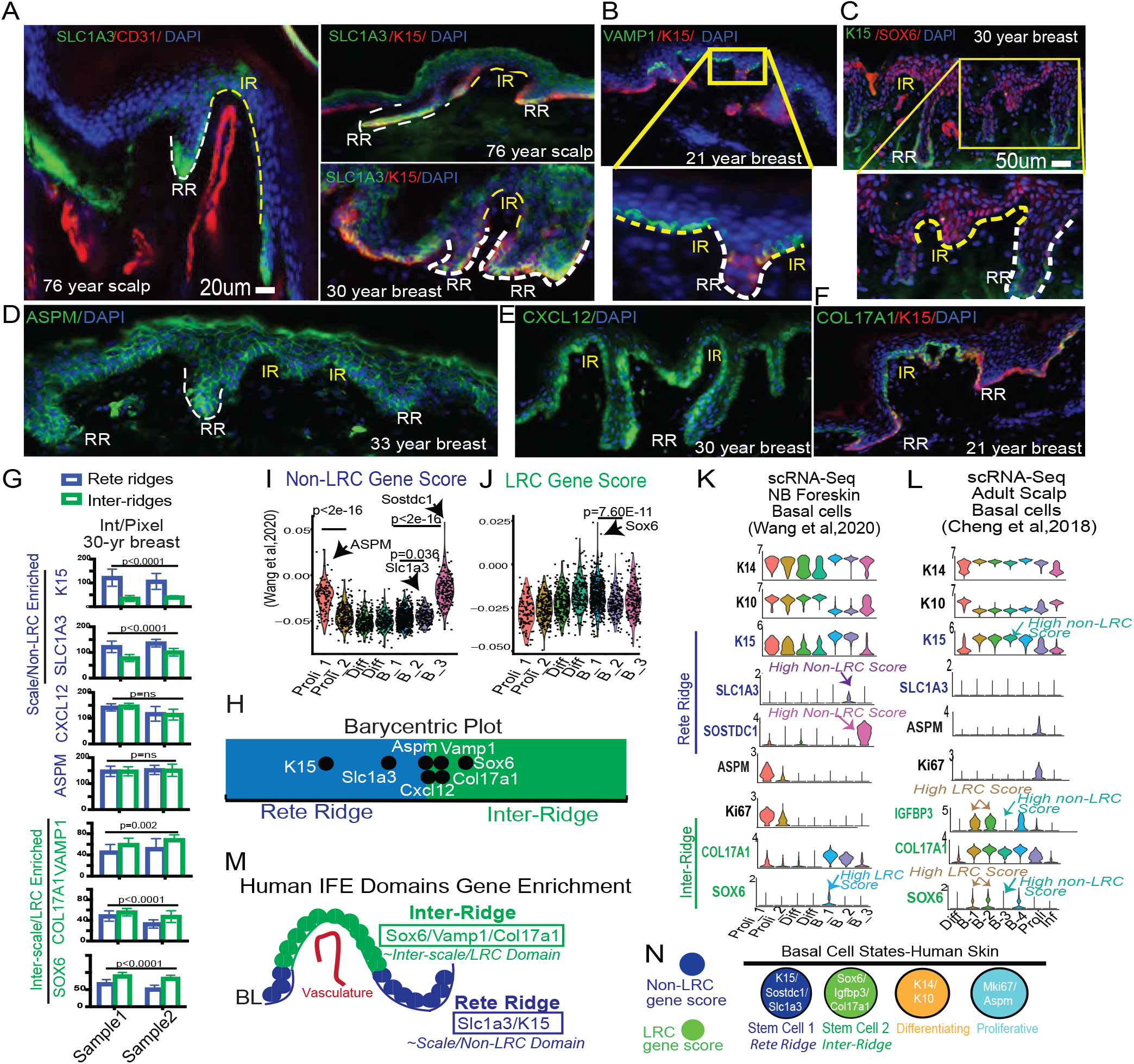
Human IFE shows basal domains and cell clusters enriched in mouse LRC/non-LRC domain markers. (A-F) Human skin immunofluorescence images of mouse basal non-LRC/LRC marker staining in rete ridges (RR) versus inter-ridges (IR). Representative images from 30 samples of different body regions and ages (see Table S1). (G) LRC non-LRC marker expression in RR and IR samples from two women, quantified using background subtraction (Material and Methods). P-values from paired Student’s t-test. (H) Relative RR vs IR gene expression/area normalized to sum to one, shown as barycentric plot. A gene expressed only in inter-ridges would be plotted at 1. Error bars are SEMs. (I,J) Gene score analysis in cell clusters from (Wang *et al*., 2020). P values from pairwise Wilcoxon rank-sum tests with Benjamini-Hochberg correction. (K,L) scRNA-seq identified clusters show specific gene expression of relevant markers. (M) Cartoon summarizing differential RR vs IR marker distribution. (N) Model of human IFE basal cell states with LRC/ non-LRC gene expression distributions

To further investigate our mouse IFE LRC/non-LRC markers in human skin, we used Seurat analysis (Butler *et al*., 2018; Satija *et al*., 2015) with Uniform Manifold Approximation and Projection (UMAP) (Korsunsky *et al*., 2019; McInnes *et al*., 2018) to probe two human single cell (sc)RNA-seq databases recently generated from newborn foreskin (Wang *et al*., 2020) and adult scalp (Cheng *et al*., 2018). We identified 7 clusters in human basal layer IFE (Figure S4E, G), and examined marker expression of known SC, proliferation, differentiation, and our mouse LRC/non-LRC genes (Figure 2I-L and S4I, J). All clusters expressed K14, indicative of their basal layer identity, but also had some low levels of K10, suggesting they are initiating differentiation, as was also reported for mouse basal IFE (Cockburn *et al*., 2021; Lin *et al*., 2020; Park *et al*., 2017; Rompolas *et al*., 2016). Several clusters expressed Ki67, indicative of actively proliferating cells, and interestingly, those clusters also expressed our non-LRC marker Aspm. The remaining basal clusters that qualify as un-differentiated and non-proliferative (e.g. putative G0 SC states) expressed preferentially either non-LRC/scale makers (Slc1a3, Sostdc1, K15) or LRC markers (Sox6, Igfbp3, Col17a1). Furthermore, these clusters were enriched in either LRC or non-LRC computed gene scores (see Material and Methods and Table S2) (Figure 2I, J and S4I, J). We conclude that human IFE contains multiple basal cell states: differentiating, proliferative, and several putative SC states (Figure 2N). The SC states are enriched in either non-LRC or LRC gene, and their specific marker expressions place them in either rete ridges or inter-ridges (Figure 2N). This analysis adds to work and interpretation from (Wang *et al*., 2020). Importantly, the non-LRC vs LRC marker enrichment in human rete ridges vs inter-ridges resembles mouse tail scales vs interscales, respectively (compare Figures 2M and 1G and see Synopsis). This suggests an unexpected and novel link between the organization of these two tissue types.

### Single cell RNA-seq of basal IFE LRCs/non-LRCs sorted from mouse back and tail skin

To characterize the cell states present in our IFE LRC/non-LRC domains, we next used mouse tail and back skin isolated cellular subsets and performed scRNA-seq analysis. We FACS purified basal (Sca1+/α6-integrin+) IFE cells as H2BGFP LRCs, mid-LRCs, and non-LRCs from K5-tTA x pTRE-H2BGFP mice (Tumbar *et al*., 2004) (Figure 3A) and used 10X Genomics scRNA-seq technology (see Material and Methods). After quality control and filtering, we obtained a total of 13484 cells from back skin of 2 mice, or ∼ 4500 cells each for LRC, mid-LRC and non-LRC sorted population (Figure S5A-E). LRC vs non-LRC gene scores computed from previous microarrays (Material and Methods, (Sada *et al*., 2016)) confirmed the expected marker gene enrichment in our LRC vs non-LRC scRNA-seq databases (Figure 3B). Some LRC/non-LRC gene expression differences previously found by microarrays (Sada *et al*., 2016) were detectable by scRNA-seq (e.g. Igfbp3, Chit1, Sostdc1), but many others were not (Figure S5F, G). Quantitative reverse transcriptase (QRT)-PCR confirmed the microarray results for some of the individually tested genes (Figure S5H), underscoring detection limitations of scRNA-seq data. Despite these limitations, cluster analysis of scRNA-seq data helped define the basic cell states present in basal sorted LRC, mid-LRC, non-LRC IFE populations. Seurat and *Harmony* integration (Korsunsky *et al*., 2019) with quality control parameters and Uniform Manifold Approximation and Projection (UMAP) (McInnes *et al*., 2018) reduction method generated 10 basal cell clusters in back skin, well-correlated in two mouse samples (Figure S5C-E). The clusters, representing basic cell states of IFE basal layer, were present to some extent in all LRCs, mid-LRCs, and non-LRCs sorted IFE fractions, with a notably increased % of proliferative cells in non-LRCs (Figure 3C). The results were similar in cluster analysis of 2863 (1241 LRCs, 1292 mid-LRCs and 330 Non-LRCs) basal IFE cell fractions sorted from tail skin (Figure 3D and S6 A, B). To assign cell-cluster identity, we used previously published cluster-markers (Figure 3E, S5I and S6C and Table S2), cell cycle analysis (Figure S5M), and gene scores (Figure S5L and S6D) extracted from other previously defined IFE cellular subsets (Dekoninck *et al*., 2020; Haensel *et al*., 2020; Joost *et al*, 2016). This classified both the back and tail skin IFE clusters as: 3 putative G0 SC states well-correlated in tail and back skin (Figure S6E); actively proliferating cells; and basal-differentiating cells (Figure 3C-E). Thus, the data reveal that LRC and non-LRC basal IFE sorted subsets contain all the cellular clusters defined by scRNA-seq. Based on these data we can infer that the two spatial (LRC/non-LRC) IFE basal domains of mouse back and tail skin harbor comparable mixtures of basic basal cell states (Figure 3F).

**Figure 3:**
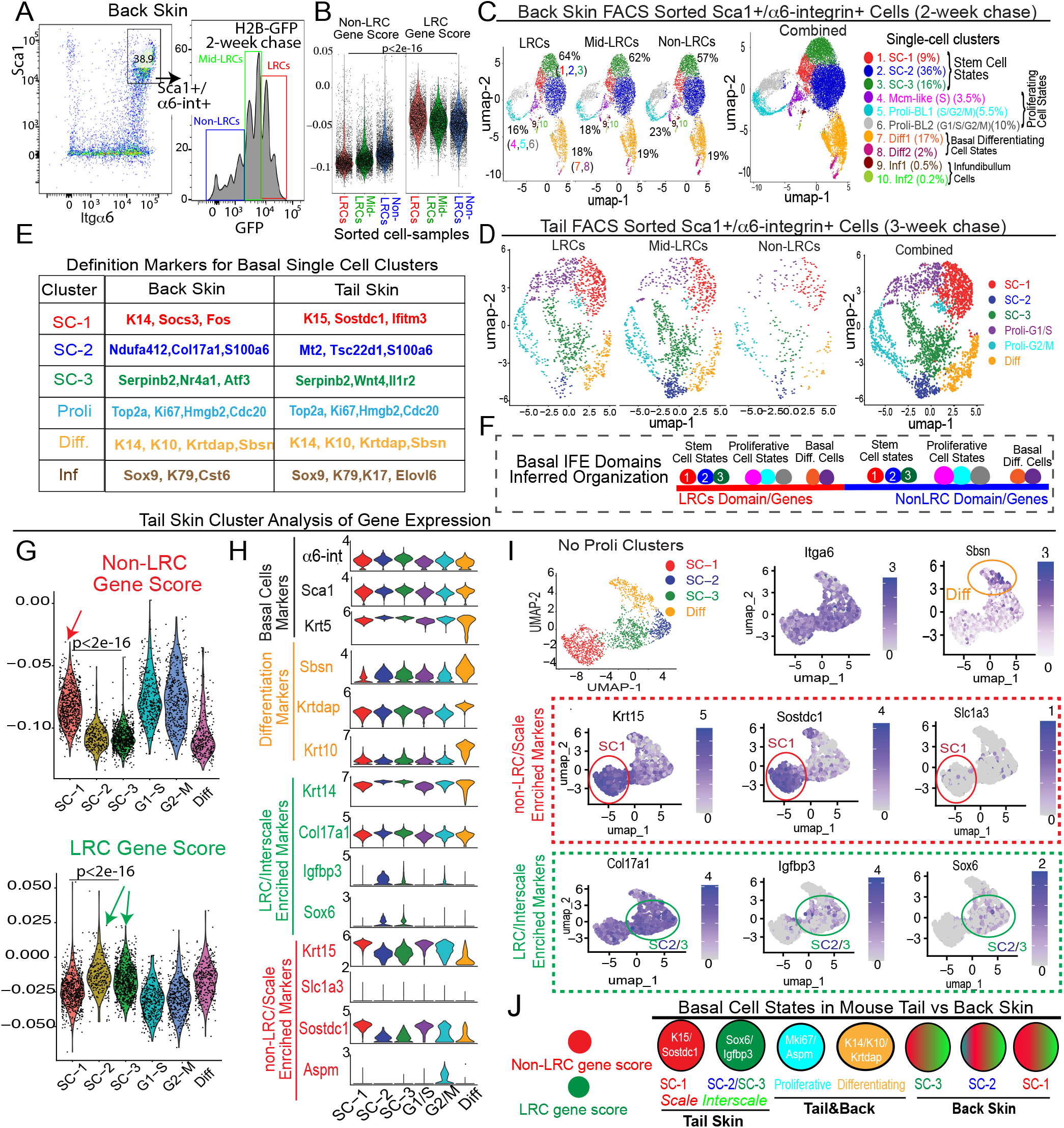
Single cell RNA-seq of basal IFE LRCs/non-LRCs sorted from mouse back and tail skin. (A) FACS sorting of basal LRCs, mid-LRCs and non-LRCs. (B) scRNA-seq data of sorted cells from (A) analyzed for LRC/non-LRC gene scores computed from microarray data (Sada *et al*., 2016). (C,D) UMAP cell clusters identified based on published markers and cell cycle regression analysis (see 3E). Marker definition analysis is described in Figure S5I-M (back) and S6C-E (tail). (E) Basal cluster definition markers. (F) Model of basal IFE domain cell states. (G) LRCs and non-LRCs computed gene scores are differentially enriched in tail SC clusters. P values from pairwise Wilcoxon rank-sum tests with Benjamini-Hochberg correction. (H,I) Violin and feature plots of markers in tail skin without proliferative (Proli) clusters. Note LRC/interscale markers Sox6&Igfbp3 enriched in SC-2&3 and non-LRC/scale markers K15/Sostdc1 enriched in SC-1. Slc1a3 is in SC-1 (scale); SC-2 expression is likely due toSlc1a3 expression in interscale line. (J) Model of LRC/non-LRC gene scores and marker expression in scRNA-seq clusters found in tail vs back skin.

Next, we examined how the LRC/non-LRC gene expression signatures defined as gene scores (Material and Methods) from the microarrays (Sada *et al*., 2016) are represented in the scRNA-seq basal cell clusters. Whereas the back skin clusters showed mild, if any, differential enrichment in LRC/non-LRC specific marker expression and computed gene scores (Figure S5J, K, N), the tail skin SC clusters showed strong differences (Figure 3G, H). Specifically, the SC-1 tail cluster was enriched in non-LRC gene scores and K15&Sostdc expressions (Figure 3G, H), which places it in the scale. Conversely, SC-2 and SC-3 tail clusters were enriched in LRC gene scores and Igfbp3&Sox6 (Figure 3G, H), placing them in the interscale. This was also apparent in feature plots, where Slc1a3 (scale marker) showed expression in SC-1 cluster (non-LRC enriched, scale) (Figure 3I). Aspm marked proliferative clusters of both tail and back skin (Figure 3H and S5J), as previously seen in human skin (Figure 2K, L). In summary, we propose that several basic basal cell states that include stem, proliferative and differentiating cells exist in both LRC and non-LRC domains of mouse back and tail skin. In the mouse back skin the LRC/non-LRC gene signatures do not differentiate the predicted stem cell states. However, in tail skin distinct stem cell states express preferentially either the LRC or the non-LRC gene expression signatures (Figure 3J), a situation also seen in human undifferentiated basal cell clusters (Figure 2N). This further underscore the previously unrecognized similarities between mouse tail and human skin.

### Two basal cell differentiation paths reflect the IFE spatial domain organization

To understand how the IFE basal cell states might relate to each other in transcriptomic lineage trajectory maps, we used Monocle 2 and Pseudotime analysis (Trapnell *et al*., 2014) from which the proliferative cell clusters were removed to eliminate bias due to cell cycle status. We analyzed all our mouse back and tail databases (Figure 4), along with two databases previously published from human (Cheng *et al*, 2018; Wang *et al*., 2020) (Figure S5F-H) and three from mouse skin (Dekoninck *et al*., 2020; Haensel *et al*., 2020; Joost *et al*., 2016) (Figure S6 I-K). Our data represents a unique mouse dataset combining both high sequencing depth and a large number of sorted cells. These combined qualities uncovered a previously unidentified bifurcated lineage tree for the basal IFE, with three differentiation branches (Figure 4A, D). This 3-branched lineage tree was also present in a human skin data set from the adult scalp (Figure 4G). The basal differentiating (K14+/K10+) cells were found on one branch, and the 3 predicted SC clusters were split on the other two branches (Figure 4A, D, G). The cell states align as a continuum on the trajectories in Monocle2, but Monocle3 predicts discrete steps along the differentiation paths more in line with current differentiation models (Cockburn *et al*, 2021; Lin *et al*., 2020) (Figure 4J, K).

**Figure 4:**
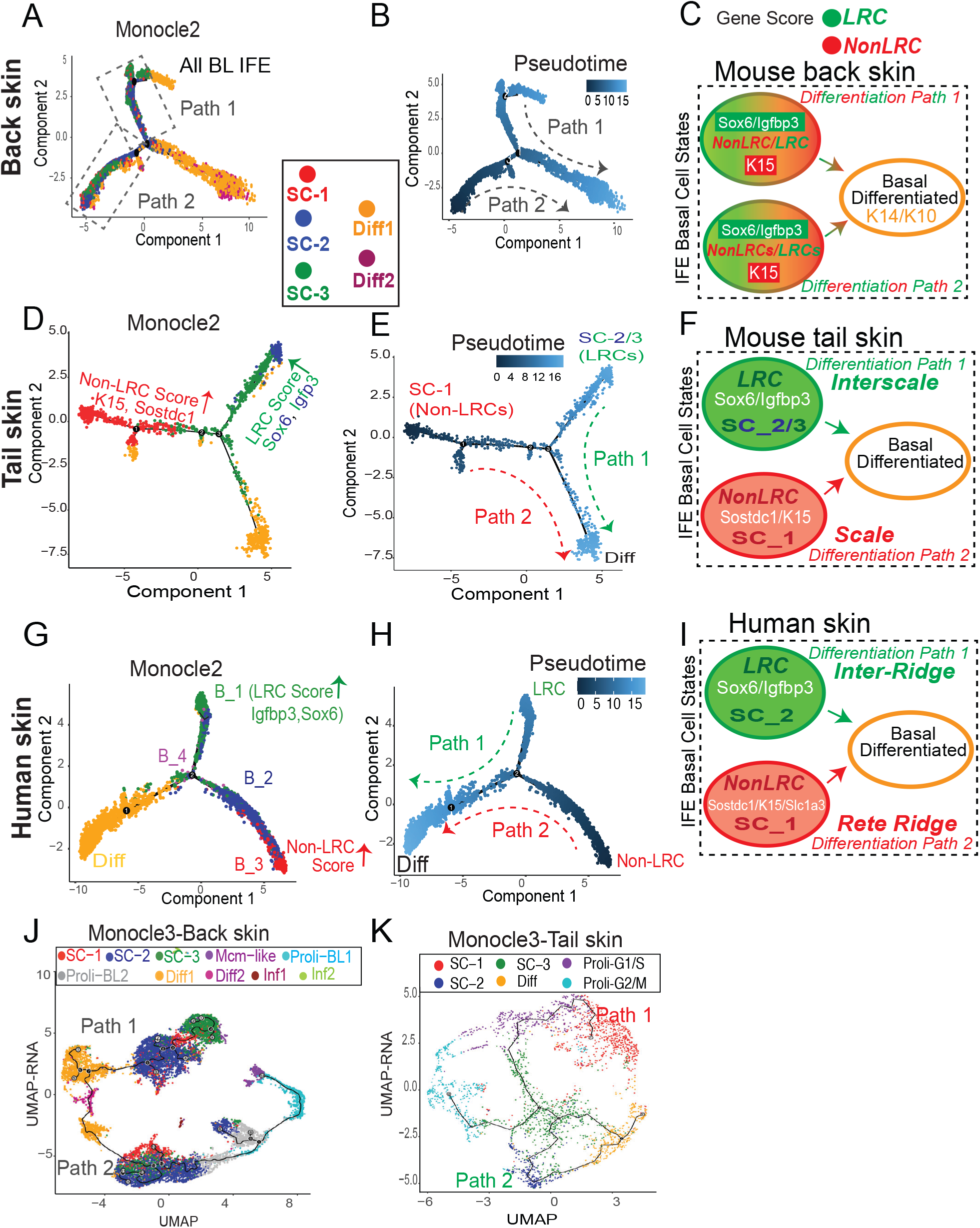
Transcriptomic Lineage Trajectory Models Reveal Two Basal IFE Differentiation Paths. (A,D,G) *Monocle 2* lineage trajectory model and associated Pseudotime predictions (B,E,H) for skin samples indicated with proliferative and infundibulum clusters excluded. (G) is human scalp from (Cheng *et al*., 2018). Pseudotime places the ‘Diff’ basal cluster branch as most differentiated, as expected based on marker expression. (C,F,I) Cartoons depicting two differentiation paths that based on LRC/non-LRC marker expression can be defined as tail scale vs interscales (F) and human rete ridges inter-ridges (I); note mixed expressions along the two paths for back skin (C). (J,K) Monocle 3 analysis of all clusters show discrete differentiation step along the two distinct paths.

Pseudotime analysis of Monocle 2 lineages assigns the two SC branches as ground states, and they both converge onto the basal differentiating cells (Diff) (Figure 4B, E, H). This suggests the existence of two distinct paths of IFE basal cell differentiation in all mouse and human tissues analyzed. In mouse back skin where LRC/non-LRC gene scores and markers were not strongly differentiated in the 3 SC states (Figure 3J), the three SC states intermingled along each of the two differentiation paths (Figure 4A-C). In contrast, in mouse tail and human scalp skin where LRC/non-LRC gene scores and markers were strongly differentiated among the stem cell clusters (Figure 3J and 2N), the SC states were strongly segregated along the two basal differentiation paths (Figure 4D, G). Specifically, in tail, SC-1 (K15/Sostdc1-enriched; non-LRC score/e.g. of the scale) was on one path, while SC-2 and SC-3 (Igfbp3/Sox6-enriched; LRC score/ e.g. of the interscale) were on the other path (Figure 4D). This distribution was similar in human skin, where a basal cluster highly enriched in LRCs score/Sox6 expression (e.g of the inter-ridge) was found on one path, while a different basal cluster enriched in non-LRC score (e.g. of the rete ridge) was found on the other (Figure 4G). Therefore, the two differentiation paths, which harbor distinct stem cell states, must reflect the spatial organization of the IFE in mouse tail scales vs interscales (Figure 4F) and in human rete ridges vs inter-ridges (Figure 4I). Two basal paths of differentiation exist in mouse back skin as well, but a possible fluidity of SC states might exist along the two paths in this distinct tissue type (Figure 4C) (see also Discussion and Synopsis).

### Molecular pathway differences suggest IFE basal domain specific adaptation to UV exposure

To further examine the cellular and molecular make up of our IFE basal populations enriched in the two spatial domains, we stained the back and tail skin of Slc1a3-CreERx, Aspm-CreERx, Dlx1-CreERxtdTomato mice at 1-month post-TM with basal markers (Slc1a3, K15, Aspm, Vamp1, and Cxcl12) (Figures 5A, B and S7A-D). K14-CreERxtdTomato mice served as a control population previously defined as a broad epidermal stem cell (Mascre *et al*., 2012), which showed even distribution of all our five tested basal markers (Figure 5B). The Dlx1-CreER, Slc1a3-CreER, and Aspm-CreER progenies were enriched in their respective ‘parent’ lineage-marker, and each showed a unique pattern of different marker distributions (Figure 5B). Notably, by 1-month post-TM, only a fraction of progenies expressed the parent-marker, suggesting that some descending cells convert into other basal cell states. This was true in both scales and interscales (Figure S7B). Therefore, we conclude that progenies of our IFE populations evolve over time, but overall remain different from each other, attesting to their distinct and heterogeneous molecular nature in the two spatial domains.

**Figure 5:**
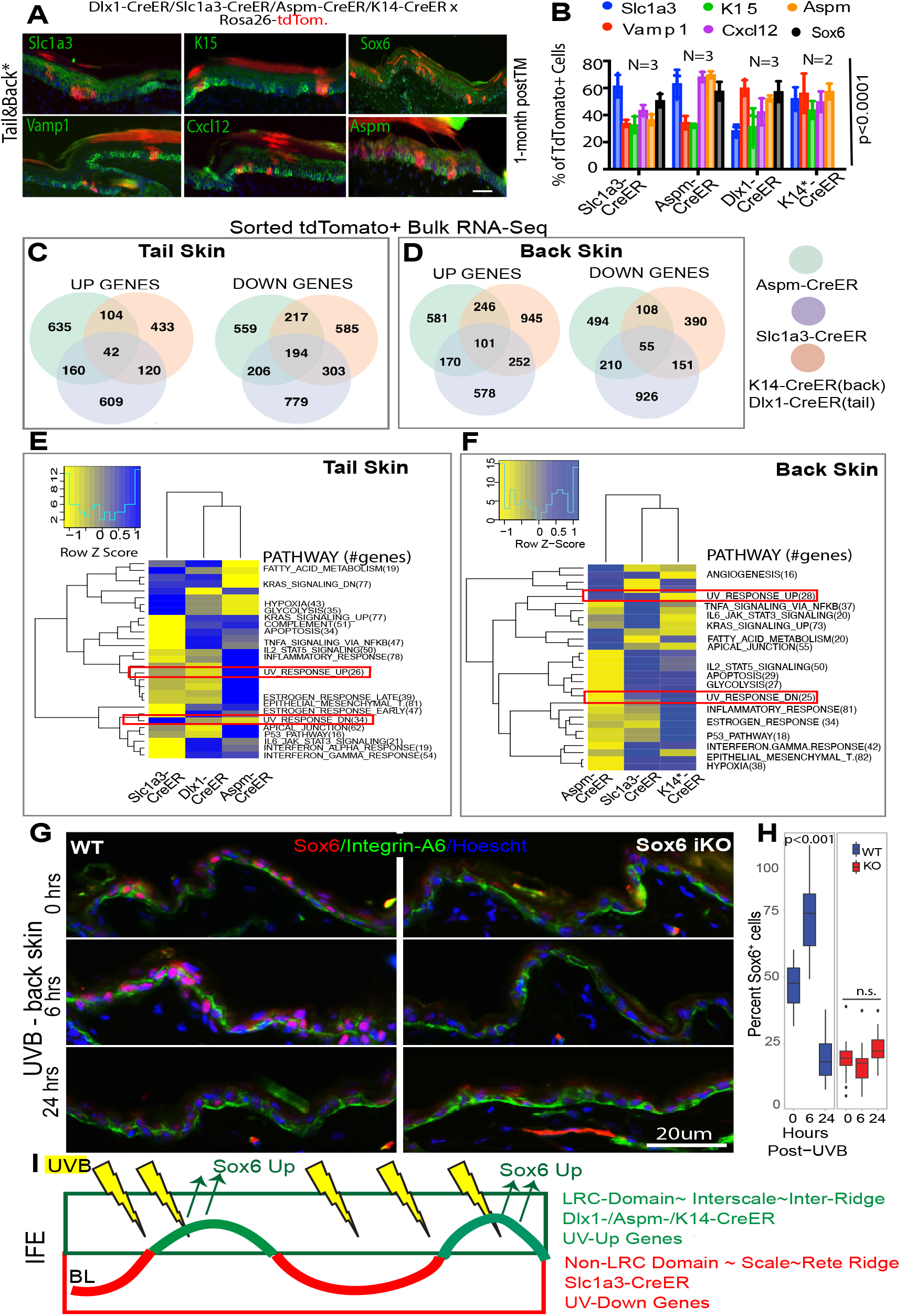
Molecular differences in IFE populations suggest domain-specific adaptation to environmental UV exposure. (A) Co-expression of markers with the lineage-traced cells at 1-month post-tamoxifen. Dlx1-CrER is shown as example; see also S9A. (B) Quantification of images like those in (A) and S9A, showing % tdTomato+ cells expressing indicated markers. Sox6 (black bar) was not analyzed in K14-Cre/EYFP lineage. See also S9B for scale vs. interscale split. (C,D) Venn diagram showing bulk RNA-seq data in the three sorted IFE basal lineages and (E,F) heatmap for their differentially regulated pathways. (G) Immunofluorescence of Sox6 staining in UVB-exposed mouse back skin. (H) Boxplot shows % Sox6 expressing basal cells quantified from images like those in (G). A linear mixed model was used for statistical analysis with 18 images from 3 mice per timepoint per genotype. (I) Model of IFE LRC/non-LRC domains and marked lineages with distinct degree of UV exposure in normal conditions.

To systematically characterize the molecular differences among these IFE populations and obtain clues on their potential physiological purpose, we turned to bulk RNA-seq analysis using 10x Illumina sequencing. For this we FACS purified >10,000 tdTomato+/Sca1+/α6-int+ (basal) cells from the back and tail skin at 2-weeks post TM induction from each of the four mouse lines. Hierarchical clustering and principal component analysis revealed hundreds of genes with differential expression in our IFE populations in both back and tail skin (Figure 5C-F and data not shown) (Table S3). GSEA analysis of hallmark pathways (Subramanian *et al*, 2005) revealed differential upregulation of many pathways including metabolism, inflammation and hypoxia (Table S4) in both back and tail (Figure 5E, F). Two pathways that differed among our sorted IFE cells especially caught our attention, as they were related to UV-response genes (Figure 5E, F). This was supported by the scRNA-Seq data, where the SC-3 population enriched in Sox6 (an interscale/inter-ridge marker) also up-regulated UV-response genes (Figure S7E). Notably, UV-induced genes such as Igfbp2, Hmga2, Col5a1and Sfrp1 extracted from previously published work (Table S2) (Li *et al*, 2021; Shen *et al*, 2019) were also found enriched in Sox6 expressing scRNA-seq clusters (e.g. SC2/3 in tail and B-1/2/4 in human scalp) (Figure S7F). That UV-pathways might be differentially regulated in the distinct IFE population and domains was especially intriguing, given that in human skin, these domains (rete ridges and inter-ridges) clearly face unequal UV exposure. Specifically, rete ridges, enriched in Slc1a3 and other non-LRC genes (Figure 2), are deeply embedded in the human skin and hence more UV-protected. Conversely, inter-ridges, enriched in LRC markers Sox6 and Vamp1 (Figure 2), are highly exposed to UV. In mouse tail, scales may be less UV-exposed than interscales, due to IFE undulations and very thick cornified envelope that retains nuclei in scales (for an example see Figure S1C). The degree of UV exposure of an IFE basal domain might potentially reflect a marker’s upregulation in that region.

To directly test if UV exposure affects expression of IFE domain markers, we irradiated the mouse back skin with UVB (Moon *et al*., 2017; Roy *et al*., 2020), which superficially affects the epidermis without penetrating to the dermal cells. We then performed IF staining for Aspm, Slc1a3, and Sox6 at several time points (Figure 5G and S7G), but only Sox6 IF signal was clearly increased in basal cells by 6-hours post-UVB exposure; no signal was detected in Sox6 KO skin, confirming Sox6 antibody specificity (Fig 5G, H). Thus, increased UV exposure of inter-ridges and interscales might explain why these IFE regions express more Sox6, and possibly other LRC-domain genes (Figure 5I). This finding poses the question of what function might LRC-domain genes like Sox6 play in skin homeostasis and in response to environmental challenges, such as UV exposure.

### Sox6 regulates IFE proliferation and survival in homeostasis and UV exposure

To first examine the role of Sox6 in the IFE during homeostasis, we deleted Sox6 from the skin epithelium and examined basal layer cell proliferation, differentiation, and survival in mouse tail. We used inducible Sox6 knockout (iKO) mice obtained by crossing the K14-CreER^T2^ (Indra, 1999) and Sox6^fl/fl^ mice (Dumitriu *et al*, 2006), and injected TM at PD32 followed by a short BrdU pulse-chase experiment (Figure 6A). We then sacrificed both wild type (WT) and iKO littermates at 2-hr, 12-hr, and 7-days of chase and quantified BrdU^+^ cells. Initially, the number of BrdU-labeled basal cells were comparable in WT vs Sox6 iKO, but by 7-day chase they significantly decreased in the iKO (Figure 6B, C, and S8A). The amount of DNA damage (Figure S8B, C), as well as the epidermis thickness marked by K10 (Figure S8D), were not significantly different between two groups. To check whether the BrdU+ cell loss in the iKO IFE during chase is caused by changes in proliferation, cell death, or both, we performed Ki67 and Caspase3 staining and quantified the results. We observed decreased proliferation (Figure 6D, E) and a transient increase in cell death in the iKO IFE compared to CT skin (Figure 6F, G). Importantly, these data indicate that Sox6 promotes cell proliferation and survival in the IFE basal layer. To our surprise, despite Sox6 upregulation in interscale compared to scale, the gene loss affected basal cells in both scale and interscale (Figure S8E-M). Apparently, even at lower expression levels as those found in the scales, Sox6 actively promotes normal IFE proliferation and survival during skin homeostasis.

**Figure 6:**
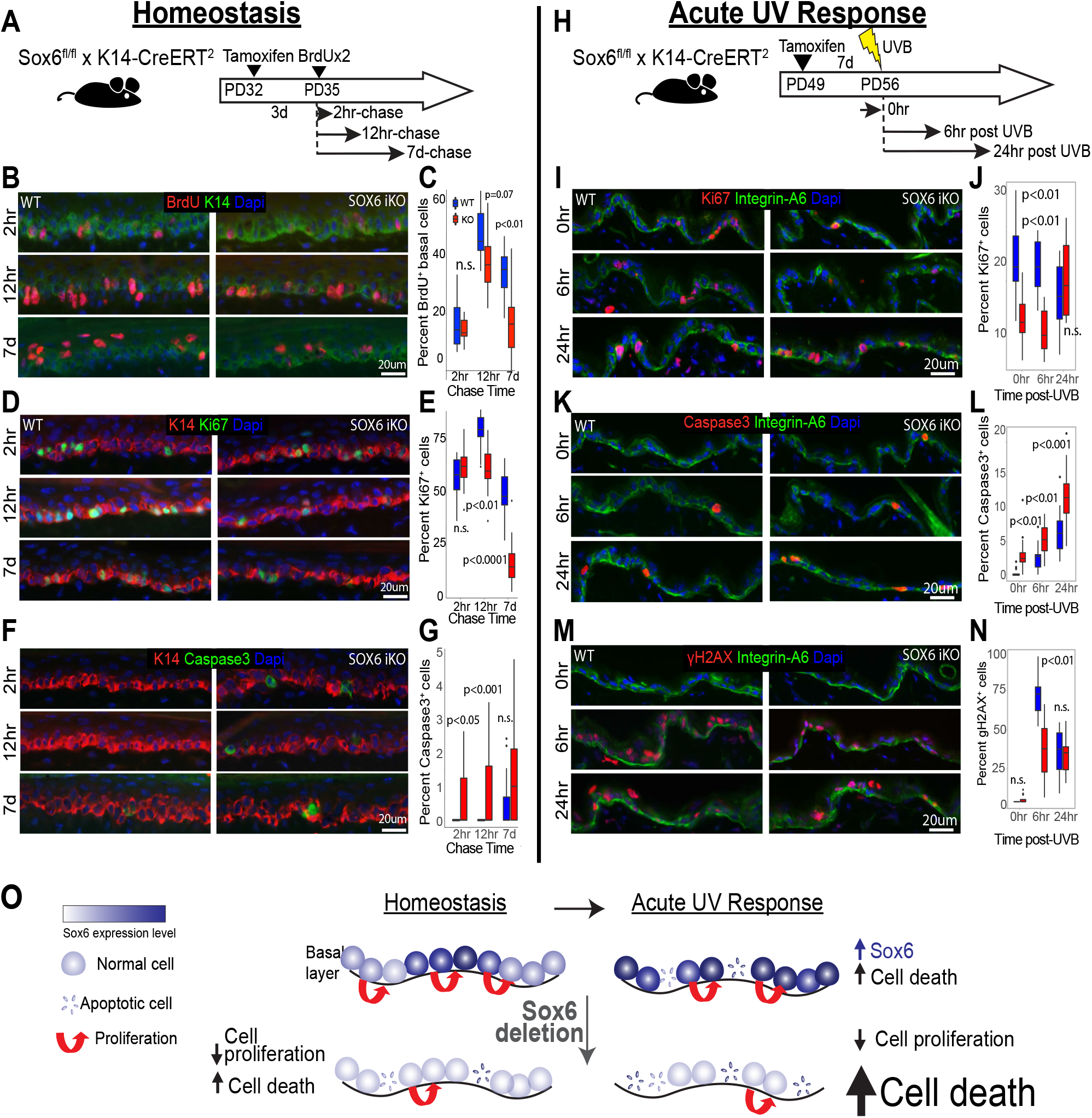
Sox6 role in basal cell proliferation and survival during homeostasis and acute UV response. (A) Schematic of Sox6 epithelial knockout (iKO) and BrdU pulse-chase experiment. (B,D,F) Images of tail skin immune-stained for markers indicated and (C,E,G) corresponding quantification box plots. (H) Sox6 role in acute UV-response was assessed in back skin to avoid compounding effects from the nucleus-retaining cornified envelope in tail scales. (I,K,M) Images of skin immune-stained for markers indicated and (J,L,N) corresponding quantification box plots. Linear mixed model was used for statistical analysis with 18 images from n=3 mice per timepoint per genotype for all quantified data in this figure. (O) Model depicting the role of Sox6 in IFE homeostasis and acute UV response.

Since Sox6 was upregulated in the IFE upon acute UVB irradiation (Figure 5G, H), we then asked if Sox6 plays a role in UV response. Thus, we UVB irradiated back skin of WT and Sox6 iKO mice and analyzed them at different time points post-exposure (Figure 6H). Staining with Ki67^+^ demonstrated that proliferative cells were fewer in the IFE iKO mice at 0 and 6-hrs post-UVB, although they recovered by 24 hr (Figure 6I, J). In contrast, cell apoptosis was consistently increased in the Sox6 iKO IFE upon UVB treatment (Figure 6K, L). Furthermore, the frequency of DNA-damaged cells was decreased in the iKO IFE by 6hr post-UV (Figure 6M, N). These results together suggest that upon Sox6 loss, IFE basal cells may resort to apoptosis rather than DNA damage repair under UVB-induced stress. Overall, our data demonstrate that the LRC-factor Sox6, enriched in more UV exposed IFE regions (e.g. inter-ridges in human and inter-scales in mouse tail), plays a vital role in basal cell proliferation and survival during homeostasis and in acute UV exposure (Figure 6O). Sox6 may represent an LRC-domain gene prototype, providing a clue with respect to one potential physiological significance of LRC vs non-LRC IFE domain organization.

## Discussion

Here we characterize the molecular, cellular, and functional organization of two types of IFE basal spatial domains in both mouse and human skin. The two spatial IFE domains were previously shown in mouse tissue kinetics studies to renew at different rates in mouse back and tail skin (Gomez *et al*., 2013; Roy *et al*., 2016; Sada *et al*., 2016), hence harboring preferentially either LRCs or non-LRCs in pulse-chase experiments. Based on lineage tracing data with different markers, we previously proposed that the two spatial domains contain distinct stem cell populations (Sada *et al*., 2016). Here we demonstrate by in situ analysis that the two basal IFE domains express LRC/non-LRC markers and some other known SC markers at different protein levels. The two domains contain comparable mixtures of molecularly discrete basal cell states on their path to differentiation. Importantly, we propose that the many-level organization of these two domains reflects in part the IFE adaptation to environmental exposure.

The spatial organization of IFE in domains is reflected in two basal cell differentiating paths, as described by our single cell transcriptomic analysis. The two differentiation paths harbor distinct stem cell states and, unexpectedly, show strong similarities in human skin and mouse tail (but not back) skin. Specifically, the tail IFE domains, defined as scales and inter-scales, show similarities of marker expression and lineage differentiation trajectories with the human rete ridges and inter-ridges respectively (Synopsis). Furthermore, our IF and single cell transcriptomic data suggest that mouse scales and their newly found human counterpart - rete ridges - harbor stem cell states enriched in non-LRC gene signatures. Conversely, the mouse inter-scales and their new human counterpart – the inter-ridges - the SCs are enriched in LRC gene signatures (Synopsis). The mouse back skin, which has so far been considered a prototype of human skin, shows the most distinct behavior among all these skin types. Although the mouse back skin presents two distinct differentiation paths, the SC states are intermingled along the paths and present a mixed identity based on LRC/non-LRC gene signatures (Synopsis). Perhaps this feature of mouse back skin data reflects the fluidity of spatial microdomain organization in this tissue. This may be influenced by the waves of synchronized hair cycle progression specific to mouse back skin (Plikus *et al*, 2011; Roy *et al*., 2016). In contrast to back skin, the two IFE domains in mouse tail (scales/interscales) and in human skin (rete ridges/inter-ridges) remain as fixed structures throughout animal’s life.

An important finding of our study is that cells from the two IFE domains express a set of genes and molecular pathways that distinguish them at the functional level, providing a link with UV-response (Roy *et al*., 2020). This may be explained by the differential exposure of the two IFE domains to the outside world (e.g. rete ridges & scales are less exposed than inter-ridges & interscales). We propose that the differential pattern of basal gene expressions in the more environment-exposed regions (e.g. the LRC gene signature) may be at least in part established because of skin’s increased UV stress in these IFE regions (Synopsis). This model is supported by our data on the LRC-factor Sox6, a novel interscale/inter-ridge enriched marker. Interestingly, Sox6 is also upregulated in skin with atopic dermatitis, a disease in which epidermal barrier defects cause excessive environmental susceptibility (Liew *et al*, 2020). Our gene targeting data in mice demonstrate that Sox6 promotes basal cell proliferation and survival during homeostasis. We also find that Sox6 is readily up-regulated by basal cells upon UV exposure to combat cellular stress and prevent excessive apoptosis (Synopsis). This response could be a broader characteristic of LRC-related genes, as some of the UV-induced master regulators (Li *et al*., 2021; Shen *et al*., 2019) appeared to be preferentially upregulated in the LRC/inter-ridge-enriched IFE domains. These observations warrant a more systematic testing of the extent of LRC signature genes implication in UV and other environmental-stress responses in the future. While exposure to UV and other environmental challenges may contribute to spatial and molecular IFE domain organization, this is likely not the origin of it. In fact, these distinct spatial IFE domains already exist to some extent in newborn human foreskin ((Wang *et al*., 2020) and this work), suggesting they may be established during skin development. We plan to investigate this fascinating question in the future.

In closing, we propose that the molecular, cellular, and spatial organization of the IFE basal layer may reflect skin adaptation to local environmental challenges, imposed on a non-uniform 3D tissue structure. This multiple-level organization may contribute to the remarkable robustness of epidermis, this extraordinarily resilient and essential tissue that faces constant challenges during adult homeostasis.

## Acknowledgements

We thank L. Tesfia and J.E. Mahoney for FACS; C. J. Bayles, R. M. Williams and J. M. DelaCruz for confocal imaging (BRC Facility) and data processing; J. Grenier, A.E. Tate and F. Ahmed for RNA sequencing (TREX facility); P.A. Schweitzer for 10x genomics scRNA-seq run; B. Cosgrove for help with scRNA-seq analysis; T.E. Hall for Vamp1 antibody staining conditions. The Cornell CARE staff for mouse husbandry. The Cornell Biotechnology Research Center and Imaging Facility as supported by NIH Grant 1S10RR025502-01. The research was supported by grants from the National Institute of Arthritis and Musculoskeletal and Skin Diseases (R01AR070157 and R01AR073806) to TT.

## Author contributions

S.G., S.L., and T.T. designed the experiments and interpreted the data. S.G., S.L., and T.T. wrote the manuscript. K.J., J.B. and S.R. performed experiments. G.C. helped with scRNA-seq analysis and trouble shooting. S.G., S.L., and D.S. performed statistical analysis and generated graphs. P.K. helped with human samples acquisition and interpretation. P.K., D.S., and G.C. assisted in manuscript assembly.

## Supplementary figure legends

**Supplementary Figure 1**. Associated with Figure 1.

**H2B-GFP pulse-chase system to mark LRC and non-LRC IFE domains in mouse tail skin from lineage traced mice**

(A) Whole mount view (top) and sections (bottom) of mouse tail skin from Dlx1-/Slc1a3-/AspmCreER x K5-tTA/pTRE-H2BGFP mice showing H2BGFP label retaining cells (LRCs) after doxycycline (doxy) chases. Scale bar=50 or 200um. (B) Cartoon showing the microscopic view for whole mount and sections. (C) Top Panels: Schematics of tet-off (left) versus tet-on (right) inductions used for comparison with (Piedrafita *et al*., 2020). Bottom Panels: Tail skin sections from K5-tta x and K14-rta x pTRE-H2BGFP mice stained with interscale (K10) marker (as indicated). ‘Mosaicism’ denotes areas of the IFE devoid of H2BGFP due to deficiency in transgene expression. High vs low exposure images demonstrate enrichment of LRCs in the interscale areas in both mouse models. Better contrast is obtained in the K5-tTa (our preferred) mouse model compared to the K14-rTa (Piedrafita *et al*., 2020) and a minimum of 2-week chase is required to note differences. (D) Tail skin sections stained for LRC/non-LRC markers as in Figure 1D, exhibit preferred overlap with H2BGFP LRC or non-LRCs IFE domains.

**Supplementary Figure S2:** Associated with Figure 1.

**H2B-GFP pulse-chase system to mark LRC and non-LRC IFE domains in mouse back skin and other body regions from lineage traced mice**

(A) Schematic of tamoxifen and doxy chase times for various body skin regions. (B) Back and ear skin sections from unchased K5-tTA x pTRE-H2BGFP mice. Mosaicism is due to deficiency in transgene expression in ear skin. (C,D) Whole mount (C) and section (D) view of back from lineage-traced mice showing 2-week chased H2B-GFP (green) post-injection with high tamoxifen (TM) dose. Scale bars, 200um or as indicated. (E) Images of skin from back, ear and paw immuno-stained for some markers from Figure 1D. Note overlap with H2BGFP LRCs vs non-LRCs domains.

**Supplementary Figure S3:** Associated with Figure 1.

**Expression of select markers in newborn versus adult mouse skin**

Immunostaining for select markers analyzed in Figure 1D shown here in mouse skin from newborn (A) vs adult back (B) and adult tail (C) skin. Markers show more uniform expression in newborn back skin.

**Supplementary Figure S4:** Associated with Figure 2.

**Human skin LRC/non-LRC marker staining and scRNA-seq analysis**

(A) Ki67 staining of human skin at various ages. (B) Quantification demonstrates preferential co-localization of K15 with Slc1a3 (non-LRC/scale marker), but not with Vamp1 (LRC/interscale marker) in human skin of all ages, except in newborn foreskin. (C,D) Examples of staining in newborn foreskin and other adult body regions suggests more pronounced heterogeneity in the adult. (E,G) Seurat generated clusters from scRNA-seq extracted from indicated studies shown as UMAP plots. (F,H) Feature plots show expression of specific markers in clusters from (E) and (G), respectively. (I,J) Non-LRC and LRC gene enrichment score analysis in cell clusters from (Cheng *et al*., 2018). P values were calculated using pairwise Wilcoxon rank-sum tests with Benjamini-Hochberg correction.

**Supplementary Figure 5:** Associated with Figures 3 and 4

**Mouse back skin scRNA-seq of sorted basal LRCs, Mid-LRCs, Non-LRCs**

(A) mRNA or gene count and percent mitochondrial genes reflecting the quality of selected cells used for scRNA-seq analysis are shown as violin plots for the (A) two mouse replicates and (B) three sorted IFE cell samples and (C) ten clusters used in our scRNA-seq experiment. (D,E) High correlation between UMAP plots (D) and clusters in gene expression matrix (E) from the two mouse replicates. (F,G,H) Gene expression changes in sorted basal cells from microarray (Sada *et al*., 2016)(F), scRNA-seq (G) and qRT-PCR (H); SEM and p values are indicated. N=3 in (F), 2 mice in (G) and 4-5 mice in (H). Note low or undetectable levels of many markers in (G). Two-tailed Student’s t-test. (I) Violin plots showing expression of genes extracted from refs (Dekoninck *et al*., 2020; Haensel *et al*., 2020) utilized to assign cell identity to each cell cluster. (J) Violin plot representation for the LRC/non-LRC marker expression in the given clusters across the samples showing generally low expression or weak differences in clusters. (K) Feature plots of cluster-defining markers and LRC/non-LRC genes. (L) Scores extracted from previous publication as indicated (Dekoninck et al., 2020; Haensel et al., 2020; Wang et al., 2020) demonstrate correlations amongst our 3 SC populations with the other databases (Table S2). P value was calculated using pairwise Wilcoxon rank-sum tests with Benjamini-Hochberg correction. (M) Cell cycle scoring of cells in various clusters as defined by cell-cycle phase-specific genelists using Addmodulescore (see methods) function in Seurat. (N) LRCs and non-LRCs gene scores from previous microarray of back skin (Sada *et al*., 2016) detect both gene sets in all single-cell derived clusters with some enrichment.

**Supplementary Figure 6:** Associated with Figure 4

**Tail and human skin scRNA-seq analysis of sorted basal LRCs, Mid-LRCs, Non-LRCs**

(A) mRNA or gene count and percent mitochondrial genes showing the quality controls for cells used for scRNA-seq analysis are shown as violin plots for the tail sorted samples and its (B) six clusters. (C) Violin plots showing expression of genes to assign cell identity to each cell cluster and (D) SC gene scores extracted from (Dekoninck *et al*., 2020). These data indicate correlations amongst the 3 SC populations from our tail databases. (E) Correlations of our tail and back skin SC clusters. (F-H) Lineage trajectory using Monocle 2 showing all clusters (left) or non-proliferative only clusters (middle) from human foreskin (Wang *et al*., 2020) and all clusters from adult human scalp (Cheng *et al*., 2018) (right). (I-K) No Proli(ferative) clusters extracted from published databases as indicated (Dekoninck *et al*., 2020; Haensel *et al*., 2020; Joost *et al*., 2016) used to generate Monocle 2 lineage trajectories.

**Supplementary Figure 7:** Associated with Figure 5

**Molecular characterization of lineage-traced IFE populations**

A) Images showing immunofluorescence staining of all the IFE markers within the lineage-traced Slc1a3-CreER, Aspm-CreER, Dlx1-CreER and K14-CreER basal cells in the tail epidermis. (B) Fraction of tdTomato+(EYFP for K14-CreER) cells positive for the respective markers in scale vs interscale for N= 2 or 3 mice per experiment (6-7 images per mouse). Scale bar∼200 um. (C) Images of back skin stained for indicated markers show co-localization in lineage-traced Slc1a3-CreER and Aspm-CreER labeled progenies. (D) Quantification of data in (C). (E) Heat map of pathways differentially expressed in scRNA-seq clustered generated from mouse tail skin shows differential regulation of UV-response genes. (F) UV gene scores extracted from previous publication (Li *et al*, 2021; Shen *et al*, 2019) demonstrate enrichment in SC 2/3 in mouse tail and B-1/2/4 in human scalp populations (Table S2). P value was calculated using pairwise Wilcoxon rank-sum tests with Benjamini-Hochberg correction. (G) Expression of Aspm and Slc1a3 in the basal layer of back skin at time points after acute UVB exposure show no apparent changes.

**Supplementary Figure 8:** Associated with Figure 6

**Sox6-loss of function phenotype in mouse skin**

(A) BrdU counts of suprabasal cells complementing data in Figure 7 B,C. (B) Staining for a DNA damage marker of skin sections from BrdU pulse-chase mice scheme shown in Figure 7A. Irradiated skin was used as positive control (CT) for the antibody staining. (C) Quantification of images like those in (B). (D) Measurements of IFE thickness based on K10^+^ stains in skin from the BrdU pulse-chase experiment at time points indicated in Figure 7A. (E) Images showing BrdU-labeled cells in scale or K10^+^ interscale, from experiment in Figure 7A. (F-M) Box plots showing percentages of markers as indicated, split per scale vs interscale from images like those in (E). Linear mixed model was used for statistical analysis with 18 images from 3 mice per timepoint per genotype for all quantified data in this figure.

## METHODS

### Mice

All mouse work was carried out according to Cornell University Institutional Animal Care and Use Committee guidelines (protocol number no. 2007-0125). To employ the H2B–GFP tet-off system, double-transgenic K5–tTA (FVB) (Diamond *et al*, 2000)/pTRE–H2B–GFP (CD1) (Tumbar *et al*., 2004) mice were used. Mice were fed with doxy chow (1 g doxy/1 kg, Bio-serv) for the indicated chase periods, starting at 1–3 months of age. Chase times are indicated for all the different experiments.

For lineage tracing, K14-CreER (CD1) (Vasioukhin *et al*, 1999), Dlx1-CreER (C57BL6) (Taniguchi *et al*, 2011) (The Jackson Laboratory, no. 014551) Slc1a3-CreER (C57BL6) (Nathans, 2010) (The Jackson Laboratory, no. 012586) mice or Aspm-CreER (Madisen *et al*, 2010; Marinaro *et al*, 2011), were crossed with Rosa–tdTomato reporter mice (Madisen *et al*., 2010) (The Jackson Laboratory). CreER/Rosa–tdTomato mice without TM injections were used to examine the leakiness of Cre. Dlx1-CreER, Slc1a3-CreER or Aspm-CreER quadruple-transgenic mice (CreER/Rosa–tdTomato/K5–tTA/pTRE–H2B–GFP) were obtained after several steps of intercrossing the above lines. Cre-ER/K5tTa (using Tet860-1060 primers), Rosa-EYFP and tdTomato mice were genotyped as recommended by the Jackson manufactures primer and protocol.

For short-term BrdU pulse-chase and UVB-irradiation experiment, K14-CreER^T2^ mice (Indra, 1999) were crossed with Sox6^fl/fl^ mice(Dumitriu *et al*., 2006) to generate an inducible epithelial-specific Sox6 KO mice.

### Tamoxifen injection

K14-CreER-H2BGFP quadruple-transgenic mice were injected intraperitoneally with a single dose of TM (Sigma) (75 μg g−1 body weight for FACS, 200 μg g−1 in K14-CreER-EYFP). For lineage tracing and FACS experiments (using Dlx1-CreER and Slc1a3-CreER lines), mice were injected with TM (100 μg g−1 body weight) for five consecutive days at 4–7 weeks of age.

Clonal lineage tracing dose was one single injection of 100 μg g−1 body weight for Dlx1-CreER and 10 μg g−1 body weight for Slc1a3-CreER or Aspm-CreER. Mice were euthanized at the indicated times after the last injection.

To induce epithelial-specific Sox6 deletion for short-term BrdU pulse-chase and UVB experiments, K14-CreER^T2^ x Sox6^fl/fl^ mice were injected with a single dose of 100 μg g−1 body weight tamoxifen at PD32 or PD49, respectively.

### Whole-mount immunostaining in the tail epidermis

Tail skin pieces (5 mm × 5 mm) were incubated in EDTA (20 mM)/PBS on a shaker at 37 °C for 2 h to separate the epidermis from the dermis as an intact sheet. Epidermal sheets were fixed in 4% paraformaldehyde (PFA) overnight at 4 °C. The skin pieces were washed, incubated in blocking buffer (1% BSA, 2.5% donkey serum, 2.5% goat serum, 0.8% Triton in PBS) for 3 h at room temperature, and incubated with primary antibodies/blocking buffer overnight at room temperature. Samples were washed 4× in PBS with 0.2% Tween for 1 h at room temperature, and were incubated overnight with secondary antibodies at 4 °C. After washing, samples were counterstained with Hoechst for 1 h and mounted. For the H2B-GFP samples, back and tail skin were chased for 2, 3 or 6-week respectively. No-GFP staining was done to enhance the signal.

Primary antibody dilutions: mouse anti-K10 (1:100, BioLegend no. 904301), rat anti-b4-integrin (1:200, BD bioscience), guinea pig anti-K31 (1:100, PROGEN Biotechnik no. GP-hHa1), rabbit anti-K14 (anti-K14 (Covance, PRB-155P). All secondary antibodies (TxR, FITC, Cy5 or Alexa-594, Jackson ImmunoResearch) were used at a 1:500 dilution. The MOM kit (Vector Laboratories) was used for blocking when staining with mouse primary antibodies.

Preparations were analyzed by confocal microscopy (Zeiss LSM710 or Zeiss LSM880) with Zen 2012 software. All confocal data are shown as projected *Z*-stack images viewed from the basal surface.

### Pre-fixation of tdTomato expressing tissues before embedding and immunostaining of mouse and human skin sections

For lineage traced mice (both tdTomato or EYFP), back and tail skin (with intact dermis) were prefixed in 4% PFA overnight and passed through sucrose gradient (15% and 30%) before embedding. Non-CreER or only H2BGFP chased mice back and tail skin was directly embedded in Optimal Cutting Temperature (OCT) compound (Tissue Tek, Sakura). The frozen sections (10 μm) were fixed with 4% PFA for 10 min at room temperature. After blocking in normal serum, sections were incubated with primary antibodies overnight at 4 °C. The following day, the sections were washed and incubated for 1 h with secondary antibodies at room temperature. After washing, the sections were counterstained with Hoechst and mounted. For staining with anti-BrdU antibody, the sections were treated with 2 M HCl for 55 min at 37 °C after blocking and stained as described above.

Primary antibody dilutions were: rabbit anti-K14 (1:1,000, BioLegend no. 905301), mouse anti-K10 (1:100, BioLegend no. 904301), guinea pig anti-K31 (1:100, PROGEN Biotechnik no. GP-hHa1), rabbit anti-K14 (Covance, PRB-155P), chicken anti-K15 (1:150, BD Biosciences no. 553731), rabbit anti-Ki67 (1:100, Leica Biosystems no. NCL-Ki67p), rabbit anti-Slc1a3(1:300, abcam 416), Aspm (1:1,000, Proteintech AB19013), Vamp1 (1:500, Abcam no. ab24646), Col17a1 (1:20,000, Novus Biology SR46-05) and Cxcl12 (1:500, cell signaling. ab24646). Rabbit anti-Sox6 (1:4,000, Abcam ab30455), anti-Caspase3 (1:4,000, R&D Systems AF835), gamma-H2AX (1:4,000, Abcam ab2893), rat anti-α6 integrin (1:4,000, BD Biosciences 555734), anti-BrdU (1:400, Abcam AB6326), chicken anti-K14 (1:40,000, 906004 Biolegend), mouse anti-K10 (1:4,000, Abcam 9026). All secondary antibodies (TxR, FITC, Cy5 or Alexa-594, Jackson ImmunoResearch) were used at 1:500 dilution. For mouse primary antibodies, the MOM kit (Vector Laboratories) was used for blocking.

Preparations were examined using a fluorescent microscope (Nikon) and digitally imaged using a CCD (charge-coupled device) 12-bit digital camera (Retiga EXi; QImaging) and IP-Lab software (MVI).

### FACS sorting

Mouse back/tail skin was incubated in 0.25% trypsin/versene overnight at 4 °C and for 30 min at 37 °C. Single-cell suspensions were prepared by scraping off the fat and subcutaneous tissue from the dermal side of the skin followed by enzymatic digestions and subsequent filtering with strainers (70 mm, followed by 40 mm). Cells were stained with the following antibodies for 30 min on ice: Ly6A (Sca1-FITC 557405 /PE-cy7 558162) from BD Biosciences and α6-integrin–Pacific blue (1:100, BD Biosciences). Dead cells were excluded by Live/Dead dye (AmCyan) staining. FACS (FACS Aria, BD Biosciences) analyses were performed in the Cornell Flow Cytometry facility. FACS data were analyzed with the FlowJo software.

### Short-term BrdU pulse-chase

For short-term bromodeoxyuridine (BrdU) pulse-chase labeling, 35 days old K14-Cre^ERT2^/ Sox6^fl/fl^ mice and their control litter mates (n=3 per group) were injected with 50 ug/g body weight BrdU intraperitoneally twice at 12hr intervals. 2hr-chased mice were harvested 2 hours after the first BrdU injection, while 12hr- and 7d-chased mice were harvested at the indicated times after the last injection.

### UVB-irradiation

UVB irradiation was performed on 56 days old K14-CreER^T2^/Sox6^fl/fl^ mice and their control litter mates (n=3 per group) following the procedure described in a previously published protocol (Moon *et al*., 2017). Isoflurane was used to anesthetize mice during dorsal skin shaving at PD55 and UVB-irradiation procedure at PD56, and 180mJ/cm^2^ UVB was irradiated on dorsal skin during the irradiation. No UVB (0hr UVB) and UVB-exposed skin samples (6hr or 24hr post-UVB) were collected at indicated times after the procedure.

### Quantification of microscope images

Flourescence intensity per unit area/pixel or fraction of tdtomato+ cells of the basal layer were quantified by using ImageJ software. The scale/interscale regions are defined based on the retention of nuclei in the cornified layer in the scale region and/or K10 (interscale) or K31(scale) expression. The data was normalized by subtracting background intensity (the region of lowest intensity in the basal layer) per image. For whole mount clonal data, the number of tdTomato^+^ clones of the tail epidermis were counted on maximal projections *Z*-stack confocal images. Clones are defined as clusters of cells that contain at least one basal or suprabasal cell. Quantifications were independently performed on ≥2 mice/per time point/per genotype, and ≥50 clones/structure were counted per mouse.

### RNA isolation and RT–PCR

Total RNAs were isolated from sorted skin cells prepared by TriZol method and used for reverse transcription by Super script III (Invitrogen). The primers used were as follows: Gapdh, 5′-ACTGCCACCCAGAAGACTGT-3′ and 5′-GATGCAGGGATGATGTTCT-3′; Dlx1, 5′-ATGCCAGAAAGTCTCAACAGC-3′ and 5′-AACAGTGCATGGAGTAGTGCC-3′; Igfbp3, 5′-TCTAAGCGGGAGACAGAATACG-3′ and 5′-CTCTGGGACTCAGCACATTGA-3′; Sox6, 5′-GGTCATGTTTCCCACCCACAA-3′ and 5′-TTCAGAGGGGTCCAAATTCCT-3′; Slc1a3,5′-ACCAAAAGCAACGGAGAAGAG-3′ and 5′-GGCATTCCGAAACAGGTAACTC-3′; Aspm, 5′-TGGCTATGAGTGAATGCTCTTCC-3′ and 5′-TCGCGTAAAAACAGTGGCAAG 3′; Vamp1,5′-CAGTGCTGCCAAGCTAAAAA-3′and 5′-CCAGTAGCCGTCTCCATACC-3′; Cxcl12, 5′-GCGCTCTGCATCAGTGAC-3′and 5′-TTTCAGATGCTTGACGTTGG-3′ K14, 5′-AAGGTCATGGATGTGCACGAT-3′ and 5′-CAGCATGTAGCAGCTTTAGTTCTTG-3′; K10, 5′-GGAGGGTAAAATCAAGGAGTGGTA-3′ and 5′-TCAATCTGCAGCAGCACGTT-3′; K15, 5′-GGAGGTGGAAGCCGAAGTAT-3′ and 5′-GAGAGGAGACCACCATCGCC-3′ qRT– PCR for each gene is normalized to GAPDH. The relative level for each gene is set to 1 in the control population.

### Single Cell capturing, library generation and processing of scRNA-seq data

FACS collected α6-integrin+/Sca1+ cells from back skin epidermis were further purified by GFP intensity into LRCs, Mid-LRCs and Non-LRCs from 2 male mice at PD47 (telogen). Single-cell suspension of each cell type (∼4000 live cells) were processed to generate single-cell 3′ cDNA libraries using individually barcoded 10X Chromium Single Cell 3′ gel bead and library Kit v3, according to the manufacturer’s recommendations (10x Genomics). RNA from the barcoded cells was reverse transcribed, followed by amplification, shearing 50 adaptor and sample index attachment. The final libraries were quantified using Agilent Bioanalyzer high sensitivity DNA chip and sequenced using an Illumina NextSeq-500. The raw data files were demultiplexed to generate the sample-specific FASTQ files, which were aligned to the mouse reference genome (mm10-3.0.0) using the 10X Genomics Cellranger pipeline (v3.1.0) with default parameters. Approximately 270 M reads were generated for the LRCs, Mid-LRCs and Non-LRCs libraries from the back skin, with a mean number of 68000, 56000 and 42000 reads per cell. After filtering, 3674, 4308 and 5500 cells were analyzed for back skin, detecting a median of 3000 genes per cell. For tail skin, it was nearly 130 M reads for each library with a mean number of 104000, 104000 and 193000 reads per cell were obtained for the LRCs, Mid-LRCs and Non-LRCs respectively.

### Single Cell RNA-seq data analysis

The output from the Cellranger from the 10X platform consists of a matrix of raw read counts that was further analyzed in R using the Seurat package version 3.1 (Satija *et al*, 2015). High-quality cells that contain at least 200-5000 genes with a mitochondrial gene percentage under 10% were filtered. Expression value scaling and normalization, PCA and UMAP dimensionality reductions and clustering were performed using the Seurat (Butler *et al*, 2018) R package (version 3.0.1). LRCs, Mid-LRCs and Non-LRCs were integrated using harmony (version 1.0) (Korsunsky *et al*, 2019) after scaling expression values for each sample independently using Seurat, resulting a total of 13487 cells by combining 2 replicates for further analysis, which could be further splitted back into LRCs, Mid-LRCs and Non-LRCs for sample specific information. To identify cell clusters, principle component analysis (PCA) was first performed and the top 10 PCs with a 0.55 resolution were used to obtain 10 clusters. Differentially expressed genes for all the clusters were acquired using FindAllMarkers function with log2 fold change > 0.25 using the Wilcoxon Ranked Sum test. Multiple resolutions (0.3, 0.5, 0.7, 0.9) were assessed to reveal clusters with biological significance and perform marker gene discovery. Markers were then selected by setting the threshold to all genes with an adjusted p value lower than 0.05. After filtering, cells belonging to the infundibulum (Sox9^high^, Krt17^high^, Krt79^high^), differentiated cells (Krt10^high^, Krt1^high^, Sbsn^high^), putative SC cell types (SC1, SC2, and SC3; Krt14^high^ Krt15 ^high^, Col17a1^high^), proliferating cells (enriched in either S-phase MCMs and/or G2-M cell-cycle related genes were remarkably segregated; Mki67, Top2a, Cdc20) were manually assigned based on their expression of various related genes (Dekoninck *et al*., 2020; Haensel *et al*., 2020).

### Pseudotime lineage trajectory

Trajectory analysis was performed using Monocle (Trapnell *et al*, 2014). First, the Seurat object (v3) was converted to a monocle cds (cell data set) and then loaded into Monocle2/3. We used the top 2000 differentially expressed genes that were calculated using the standard Seurat workflow to assign pseudotime values to individual cells. The cells that contain lower than 200 transcripts were removed from further analysis. The trajectories were constructed with DDRTree.

### LRC/non-LRC gene score computation and cell cycle analysis

Cell-cycle phases of different clusters were predicted using Seurat’s Cell-Cycle scoring method Each cell was scored based on the expression of certain G2M and S phase markers. Any cell that did not express either G2M or S phase markers was predicted to be in G0/G1. The AddModuleScore function in the Seurat R package was used to calculate signature scores and determine if there was any significant difference between the sub-populations. This function was widely used for scoring various gene sets from previous literature for both mice (Dekoninck *et al*., 2020; Haensel *et al*., 2020) and human (Wang et al., 2020). Specific genes in each gene set are listed in Table S2. The two-sided Wilcoxon rank sum test (Ji *et al*, 2020) was used to evaluate whether there are significant differences in the computed signature scores between two groups of cells.

### Bulk RNA-sequencing

RNA-seq were performed in triplicates (BL Slc1a3/Dlx1/Aspm-tdTomato positive BL and total BL α6-integrin ^+^ /Sca1^+^ cells) isolated from three female mice at PD49 (telogen). Total RNA was isolated from sorted cell populations using Trizol (Thermo Fisher) according to the commercial protocol with the following additions: after the first phase separation, additional chloroform extraction step of the aqueous layer in Phase-lock Gel heavy tubes (Quanta Biosciences); addition of 1ul Glyco-blue (Thermo Fisher) immediately prior to isopropanol precipitation; two washes of the RNA pellet with 75% ethanol. The RNA sample quality was confirmed using a Qubit3 (RNA HS kit; Thermo Fisher) to determine concentration and with a Fragment Analyzer (Advanced Analytical) to determine RNA integrity. The PolyA+ RNA was enriched with the NEBNext Poly(A) mRNA Magnetic Isolation Module (New England Biolabs).

### Illumina Library Preparation and Sequencing

Since our samples resulted in low input RNA (<20 nt total RNA), truSeq-barcoded RNAseq libraries were generated with the Ultra II RNA Library Prep Kit (non-directional) (New England Biolabs). Each library was quantified with a Qubit 2.0 (dsDNA HS kit; Thermo Fisher) and the size distribution was determined with a Fragment Analyzer (Advanced Analytical) prior to pooling. Libraries were sequenced on an Illumina instrument (Hiseq4000). 2×150 bp HiSeq reads were generated with a depth of 20M

### Mapping the reads

*preprocessing*: reads were trimmed for low quality and adaptor sequences with TrimGalore v0.6.0 (ref 1), a wrapper for cutadapt (ref 2) and fastQC (ref 3).Parameters: -j 1 -e 0.1 --nextseq-trim=20 -O 1 -a AGATCGGAAGAGC --length 50 --fastqc ;unwanted reads were removed with STAR v 2.7.0e (ref 4). Parameters: --outReadsUnmapped Fastx *mapping*: reads were mapped to the reference genome/transcriptome (mouse/mm10) using STAR v2.7.0e (ref 4). GeneCounts *gene expression analysis*: SARTools and DESeq2 v1.26.0 were used to generate normalized counts and statistical analysis of differential gene expression (ref 5,6). Parameters: fitType parametric, cooks Cutoff TRUE, independent Filtering TRUE, alpha 0.05, pAdjustMethod BH, typeTrans VST, locfunc median

### Software References

1) TrimGalore: Felix Krueger http://www.bioinformatics.babraham.ac.uk/projects/trim_galore/

2) cutadapt: Marcel Martin https://cutadapt.readthedocs.io/en/stable/ http://journal.embnet.org/index.php/embnetjournal/article/view/200

3) fastQC: Simon Andrews http://www.bioinformatics.babraham.ac.uk/projects/fastqc/

4) STAR: Alexander Dobin https://doi.org/10.1093/bioinformatics/bts635

5) SARTools: Hugo Varet http://dx.doi.org/10.1371/journal.pone.0157022

6) DEseq2: Michael Love http://www.bioconductor.org/packages/release/bioc/html/DESeq2.html https://genomebiology.biomedcentral.com/articles/10.1186/s13059-014-0550-8

### Quality control analysis

Principal component analysis was performed on the raw counts for the different tdTomato + sorted populations normalized in both back and tail skin with their respective BL (α6−integrin/+Sca1+) sorted control and the scores were represented in a three-dimensional scatter plot (data not shown). Hierarchical clustering was performed on all the samples to reflect sample grouping between tail and back epidermal populations (data not shown). After normalization, the genes having counts with a signal value <100 or reported ‘absent’ were excluded. The remaining genes that are ≥2-fold up- or downregulated in BL relative to BL tdTomato+ lineages were selected. Signature genes were defined as those that were ≥2-fold upregulated in one lineage over its own BL. To compare the populations gene-wise, lists of ‘≥2-fold change in Aspm/Slc1a3/Dlx1/K14-CreER; tdTomato normalized over total BL (∼400 genes) were used (data not shown). For pathway comparison, lists of ‘log FC≥1 or <1 genes without any cut-off on raw count signal (∼4000 genes) was used (Table S4). The numbers of overlapping genes are shown in the Venn diagram (Figure 6c). Gene Set Enrichment analyses using Hallmark pathways ([using GSEA 4.0 with classic parameters] (Subramanian *et al*., 2005) were performed on differentially expressed genes (Table S5).

### Statistics & reproducibility

All experiments with or without quantification were independently performed at least twice with different mice and the representative data are shown. All statistical analyses were performed either using the two-tailed Student’s *t*-test (mixed Anova) or linear mixed model as indicated in each figure legend; statistical significance was defined as *P* < 0.05.

The sample size was dictated by experimental considerations and not by a statistical method. The experiments were not randomized. The investigators were not blinded to allocation and outcome assessment during experiments.

## Data availability

The scRNA-seq raw data of both the replicates reported in this paper have been deposited in the GEO database under accession code GEO: *(in process)*.

Bulk RNA sequencing data that support the findings of this study have been deposited in the Gene Expression Omnibus (GEO) under accession codes: *(in process)*. All other data supporting the findings of this study are available from the corresponding author on request.

