## Supplementary Figures for "Epidermal basal domains organization highlights skin robustness to environmental exposure"

### Supplementary Figure 1 (Associated with Figure 1)

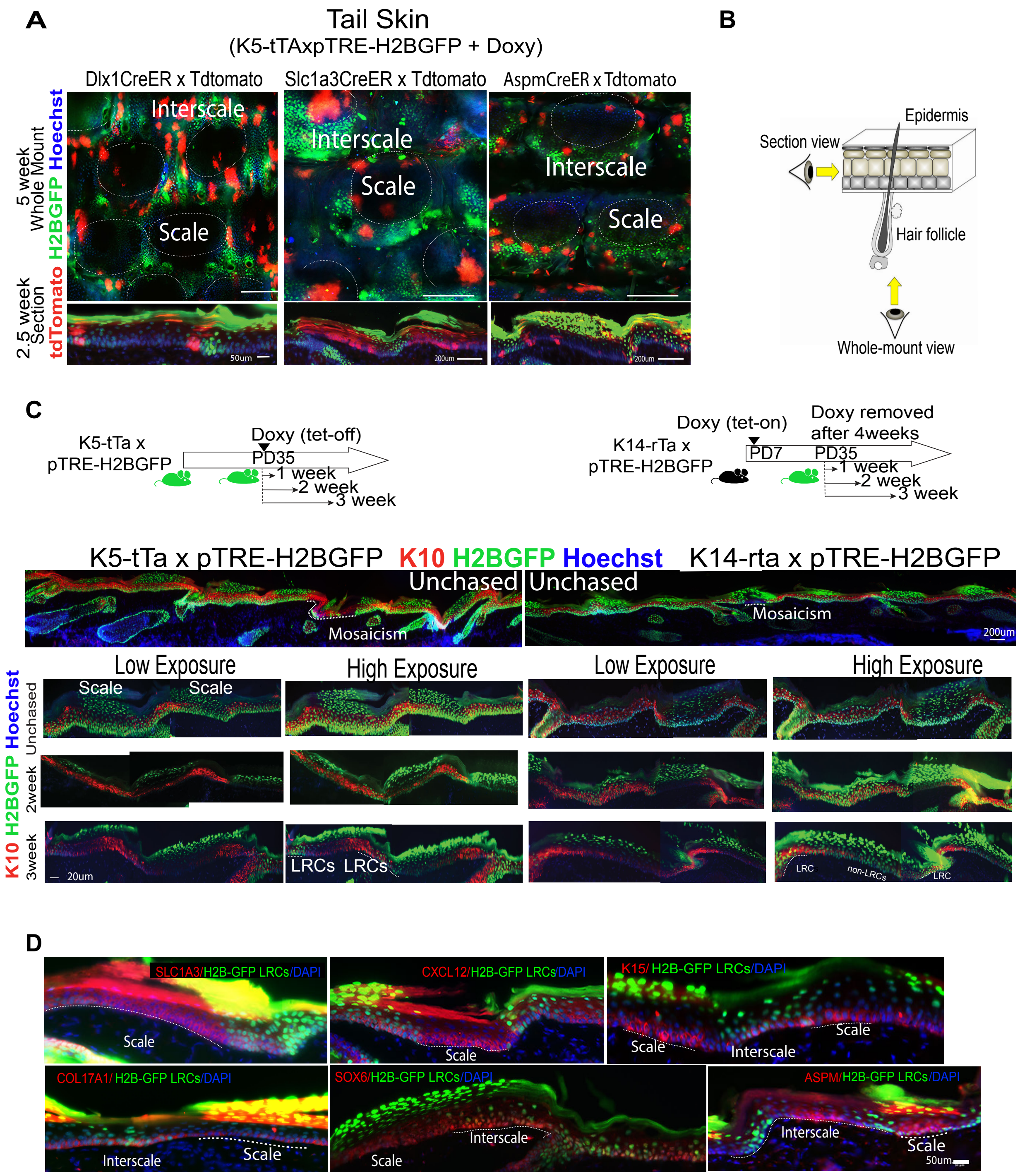

### Supplementary Figure 2 (Associated with Figure 1)

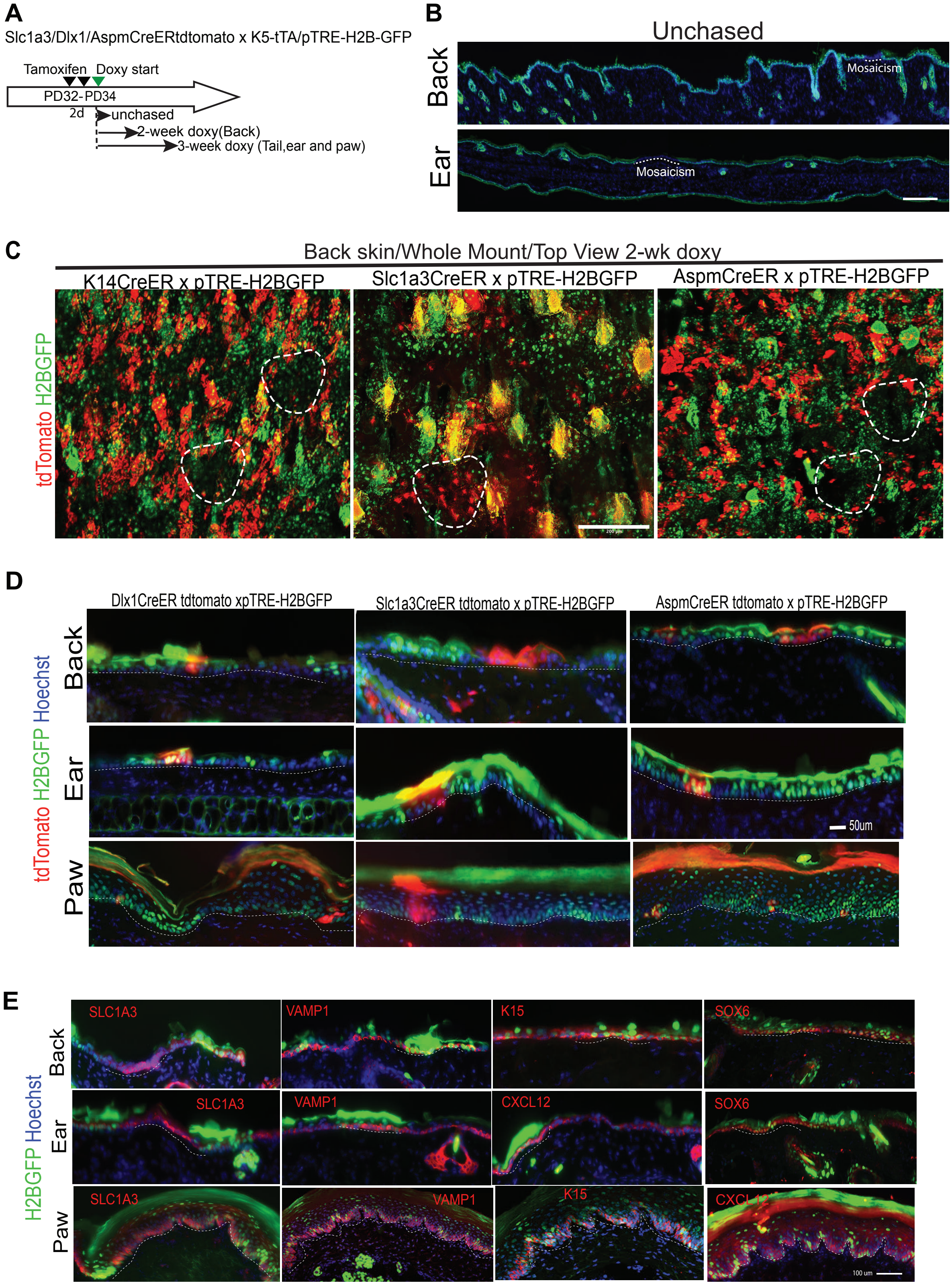

### Supplementary Figure 3 (Associated with Figure 1)

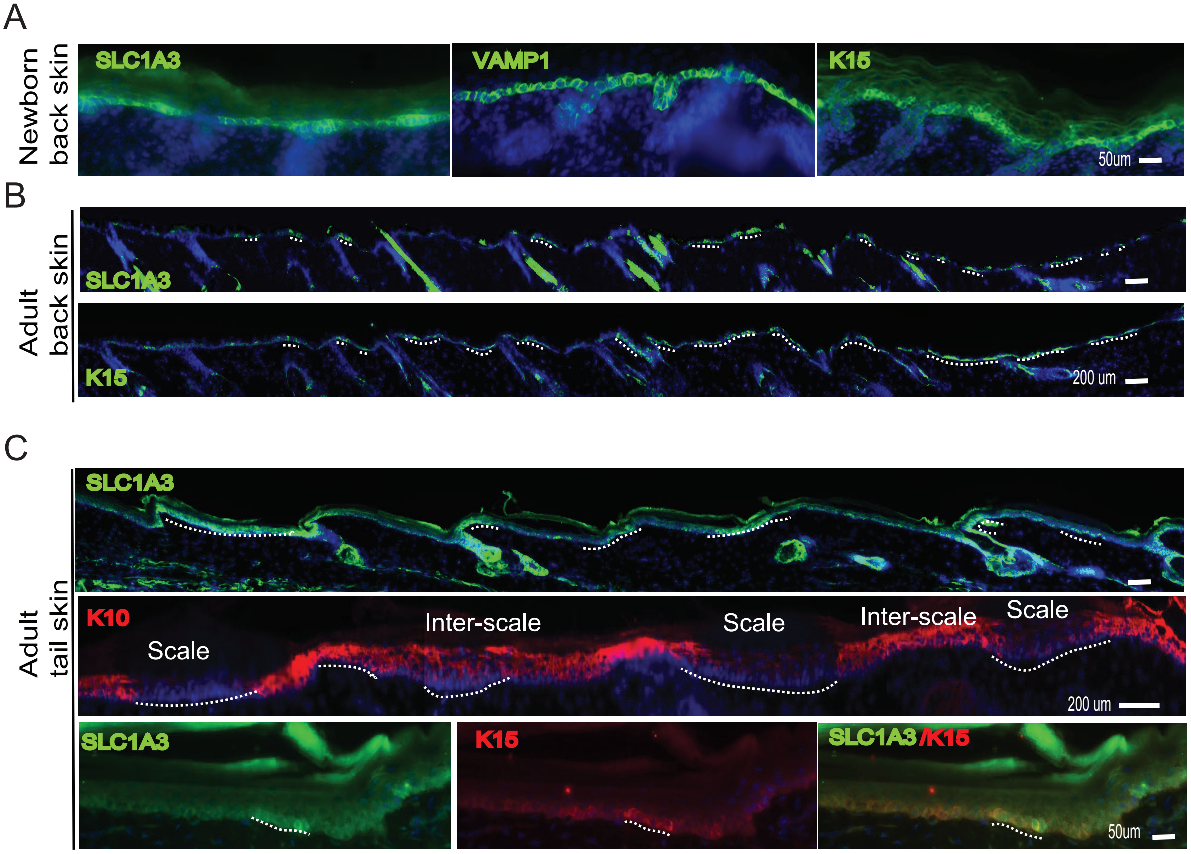

### Supplementary Figure 4 (Associated with Figure 2)

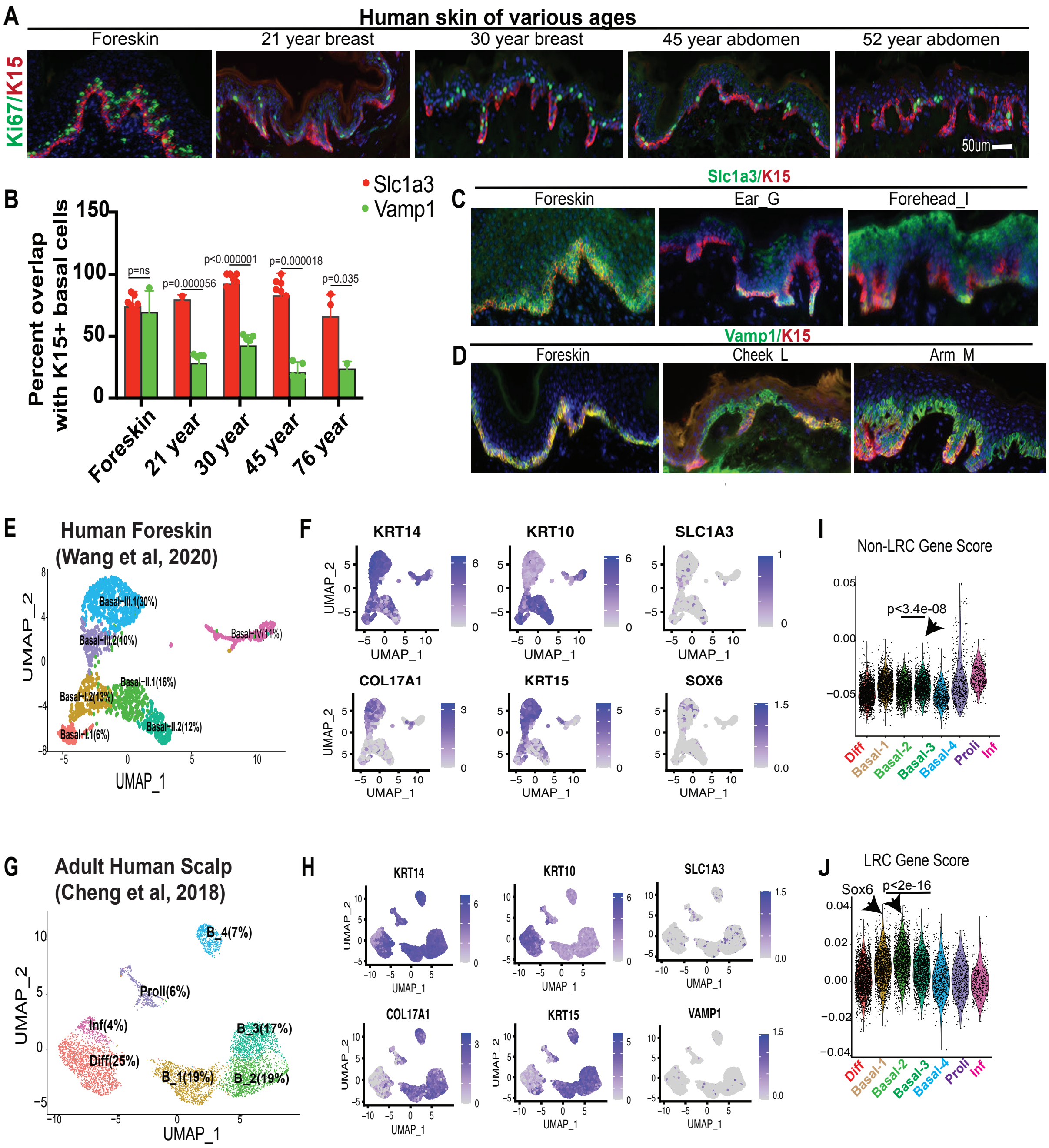

### Supplementary Figure 5 (Associated with Figure 3)

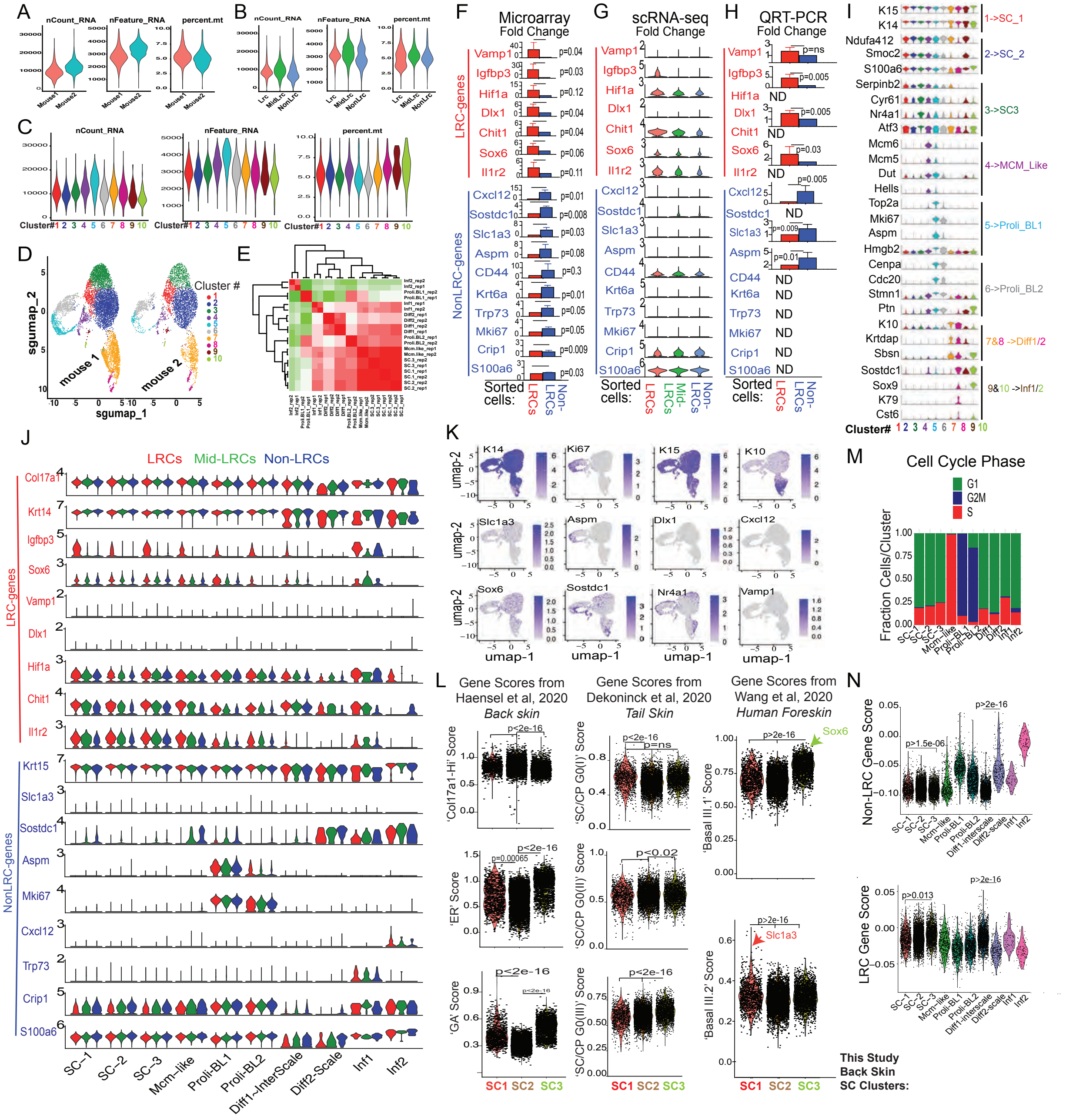

### Supplementary Figure 6 (Associated with Figure 4)

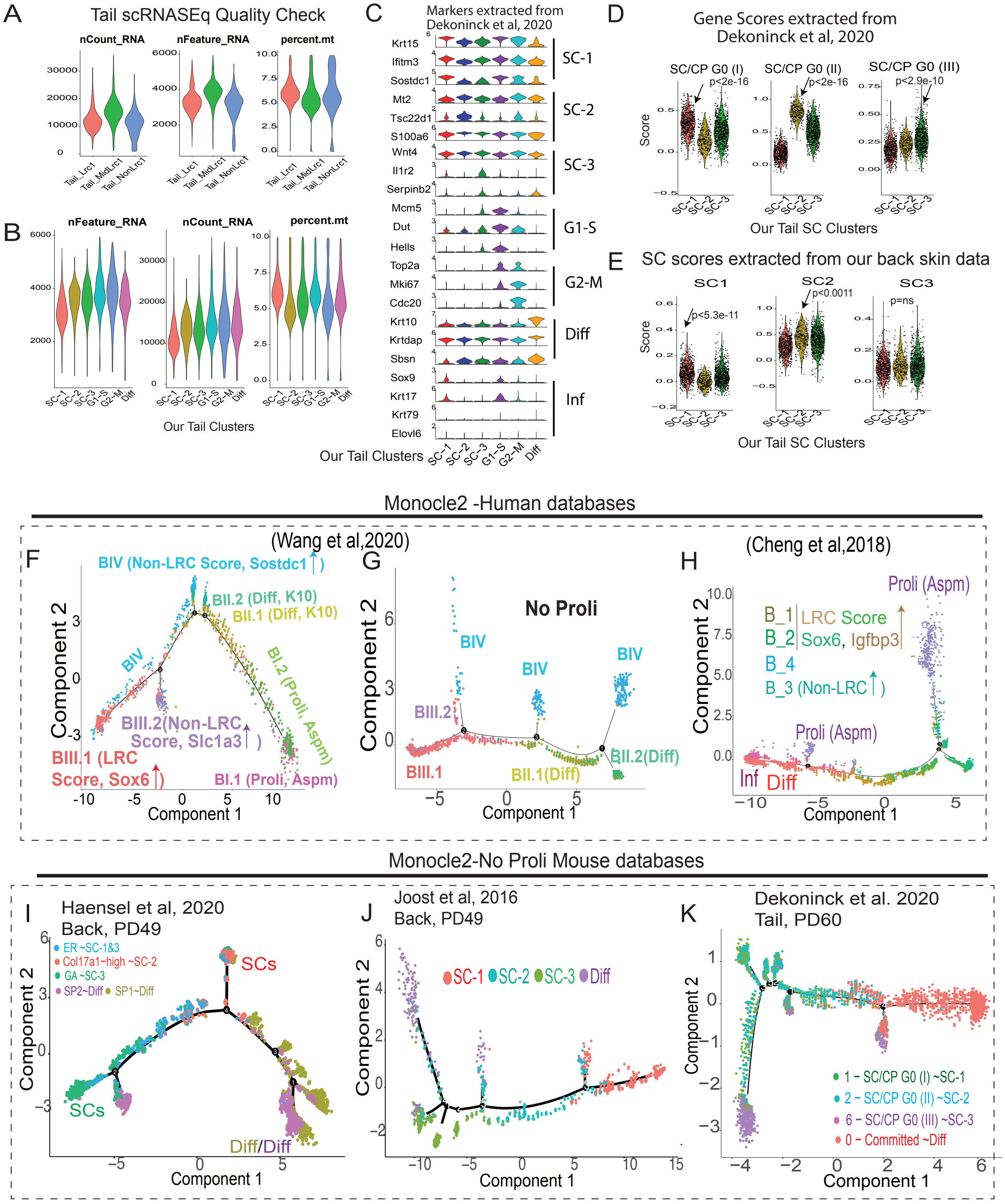

### Supplementary Figure 7 (Associated with Figure 5)

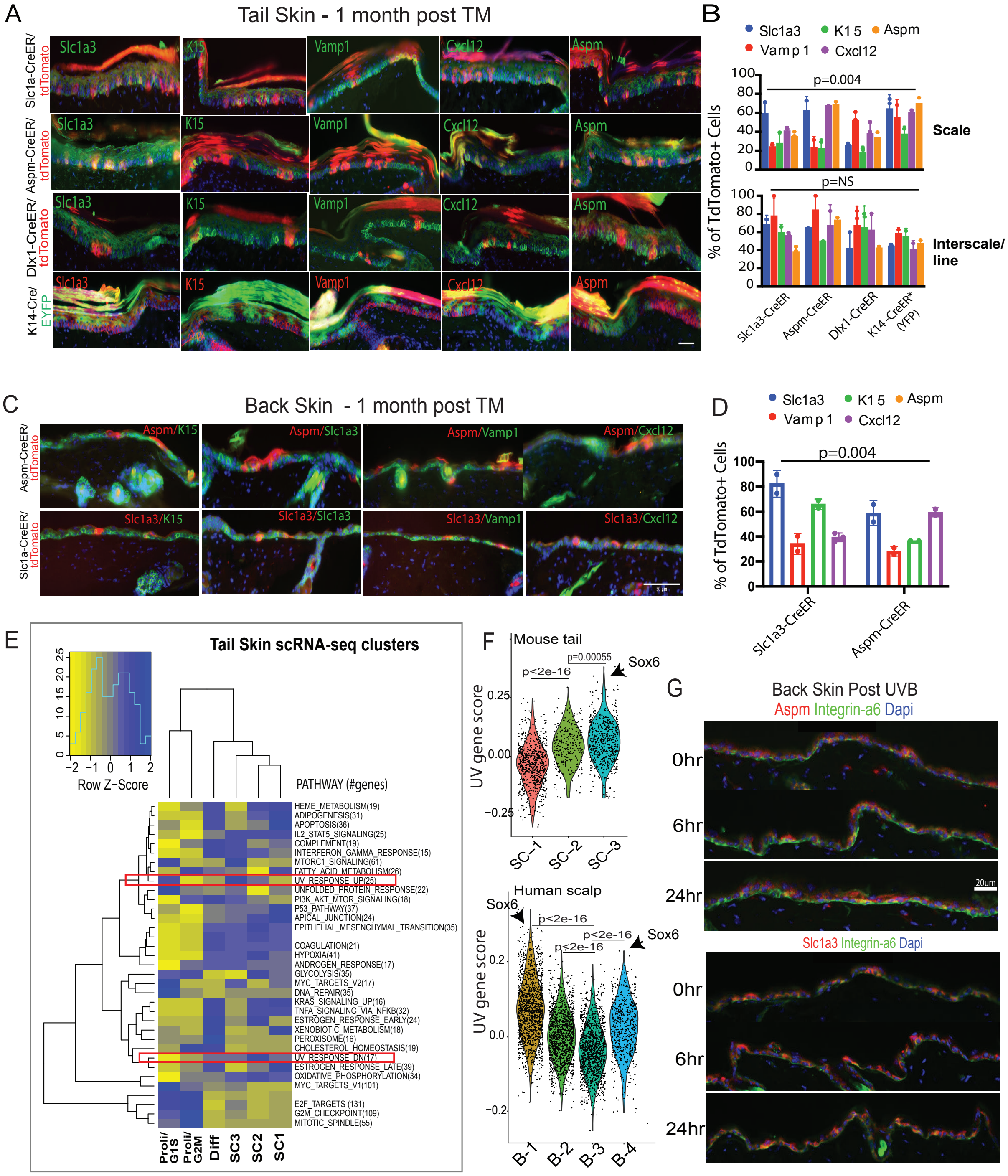

### Supplementary Figure 8 (Associated with Figure 6)

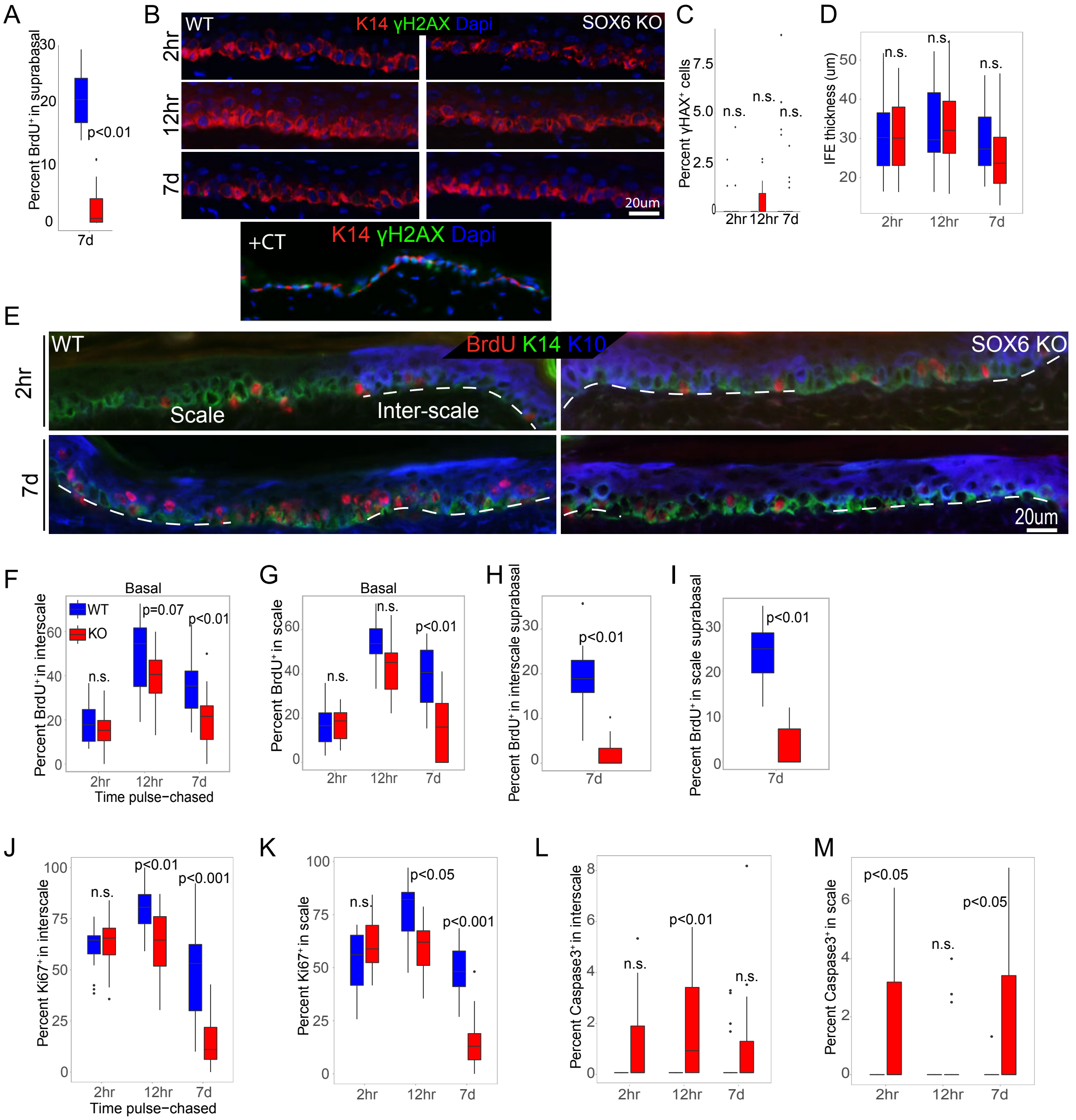
